# Transcriptional Profiling of Primate Central Nucleus of the Amygdala Neurons to Understand the Molecular Underpinnings of Early Life Anxious Temperament

**DOI:** 10.1101/808279

**Authors:** Rothem Kovner, Tade Souaiaia, Andrew S. Fox, Delores A. French, Cooper. E. Goss, Patrick H. Roseboom, Jonathan A. Oler, Marissa K. Riedel, Eva M. Fekete, Julie L. Fudge, James A. Knowles, Ned H. Kalin

## Abstract

Children exhibiting extreme anxious temperament (AT) are at an increased risk to develop anxiety and depression. Work in young rhesus monkeys mechanistically links the central nucleus of the amygdala (Ce) to AT. Here, we used laser capture microscopy and RNA sequencing in 47 young rhesus monkeys to investigate AT‘s molecular underpinnings by focusing on lateral Ce (CeL) neurons. We found 528 AT-related transcripts, including protein kinase C type-delta (PKCδ), a CeL microcircuit cell marker implicated in rodent threat processing. We characterized PKCδ neurons in the rhesus CeL, compared their distribution to the mouse, and demonstrated that a subset of these neurons project to the laterodorsal bed nucleus of the stria terminalis (BSTLd). These findings present evidence in the primate of a CeL to BSTLd circuit that maybe relevant to understanding human anxiety and points to specific molecules within this circuit that could serve as potential treatment targets for anxiety disorders.

## Introduction

Depending on its intensity and the context in which it is expressed, anxiety can be adaptive or maladaptive. Anxiety is dimensional and across a population is characterized by individual differences, and when extreme is disabling. Considerable research has identified heritable and non-heritable factors underlying the development of anxiety disorders (Eley et al., 2003; Fox et al., 2015a; Sawyers et al., 2019; Stein et al., 1999), and in childhood these can manifest as the trait-like disposition, anxious temperament (AT). Like anxiety, AT is dimensional and is characterized by individual differences in inhibitory responses to novel and social situations (Biederman et al., 2001; Davidson and Rickman, 1999; Fox et al., 2008; Hirshfeld et al., 1992) as well as threat-related pituitary-adrenal activation (Fox et al., 2008; Kagan et al., 1987). Because AT reflects a combination of behavioral and physiological responses associated with stress, this construct reflects the interplay between emotions, behavior and physiology that is emblematic of anxiety responses. When extreme and stable over time, childhood AT markedly increases the risk for the development of anxiety disorders, depression, and co-morbid substance use disorder later in life (Biederman et al., 2001; Chronis-Tuscano et al., 2009; Davidson and Rickman, 1999; Essex et al., 2010; Fox et al., 2005).

To understand the mechanisms underlying AT, we developed a young rhesus monkey model that is based on individual differences in the expression of dispositional anxiety and is analogous to the anxious phenotype exhibited by at-risk children (Fox and Kalin, 2014; Kalin and Shelton, 2003). Our approach has been to understand individual differences in the expression of the AT phenotype in relation to individual differences in its underlying neural and molecular substrates (Fox and Kalin, 2014; Fox et al., 2015a; Oler et al., 2010). Using a large multigenerational pedigree, we demonstrated that AT is approximately 30% heritable, consistent with previous studies in humans (Fox et al., 2015a; Hettema et al., 2001). Rhesus monkeys are ideally suited for studies of human psychopathology due to their recent evolutionary divergence from humans, which is reflected in similarities in brain and behavior, as well as in emotional and physiological responding (Kalin and Shelton, 2003). Additionally, the array of individual differences in emotional expression observed in humans can be readily modeled in nonhuman primates (Kalin and Shelton, 2003).

Numerous studies point to the importance of the extended amygdala in mediating adaptive responses to threat as well as stress-related psychopathology (Fox and Kalin, 2014; Fox et al., 2015b; Janak and Tye, 2015). Components of the extended amygdala include the central nucleus of the amygdala (Ce) and the bed nucleus of the stria terminalis (BST; (Alheid and Heimer, 1988). The Ce, which is primarily composed of GABAergic neurons, coordinates information that comes into and leaves the amygdala (Fadok et al., 2017; Fadok et al., 2018; Petrovich and Swanson, 1997; Veening et al., 1984; Viviani et al., 2011). In addition, the Ce sends strong projections to the BST, a region also involved in threat-responding (Dong et al., 2001; Kim et al., 2013; Oler et al., 2017). Our studies in rhesus monkeys assessing regional brain metabolism demonstrate that individual differences in Ce and BST metabolism relate to trait-like individual differences in AT and that brain metabolism in these regions is heritable (Fox et al., 2015a; Fox et al., 2008; Oler et al., 2010). When assessing co-heritability between AT and metabolism, BST threat-related metabolism was significantly co-heritable with AT whereas this was not the case for Ce metabolism (Fox et al., 2015a). Moving beyond the imaging studies, we previously used lesioning strategies to investigate the mechanisms underlying AT. Results demonstrated that selective neurotoxic lesions of the Ce reduced AT, directly implicating the Ce as a core component of the AT circuit (Fox et al., 2015a; Kalin et al., 2004; Kalin et al., 2001).

It is important to emphasize that the Ce is not uniform and can be divided into at least 2 subnuclei that include the lateral Ce (CeL) and medial Ce (CeM; (Amaral et al., 1992; De Olmos, 2004). The CeM coordinates the output of the amygdala via its projections to multiple downstream effector sites (Veening et al., 1984). The CeL modulates the CeM, helping to orchestrate the different adaptive behavioral and physiological responses mediated by the CeM’s targets (Haubensak et al., 2010; Viviani et al., 2011). The entire Ce projects to the BST to further coordinate threat-related responding, where the CeL’s projections are largely restricted to laterodorsal BST (BSTLd), (Ahrens et al., 2018; Asok et al., 2018; Oler et al., 2017; Pomrenze et al., 2019). In addition to other areas of the basal forebrain, the CeL, CeM, and the BSTLd have been conceptualized as the central extended amygdala, in contrast to what is considered the medial extended amygdala which is comprised of the medial nucleus of the amygdala and the medial divisions of the BST (Alheid and Heimer, 1988). While these subdivisions are conserved across rodent and primate species, studies elucidating the selective role of these subdivisions in mediating fear and anxiety have been predominately performed in rodents (Ciocchi et al., 2010; Tye et al., 2011). In this study, we focus on the primate CeL because studies in rodents demonstrate that it integrates information from the basolateral amygdala complex and acts as an interface between the basolateral amygdala complex and the CeM and BSTLd (Dong et al., 2001; LeDoux et al., 1990; Petrovich and Swanson, 1997; Yu et al., 2017). Studies demonstrate that within the CeL, GABAergic neuronal subtypes interact to affect CeM and BSTLd function (Ahrens et al., 2018; Asok et al., 2018; Haubensak et al., 2010; Petrovich and Swanson, 1997; Pomrenze et al., 2019). For example, protein kinase C type delta (gene name: *PRKCD,* protein name: PKCδ) and somatostatin (gene name: *SST,* protein name: SST) expressing neurons are two prominent cell types that are involved in a mutually inhibitory CeL microcircuit, integrating and gating the flow of threat-related information to the CeM and BSTLd (Fadok et al., 2018; Haubensak et al., 2010; Li et al., 2013; Ye and Veinante, 2019).

In this study, we characterized individual differences in gene expression in CeL neurons in relation to individual differences in AT and its components. Our focus on gene expression in CeL neurons was motivated by: 1) our studies demonstrating a mechanistic role for the monkey Ce, 2) the role of the CeL in gating and integrating information within the amygdala, and 3) the CeL’s role in modulating fear and anxiety-related responses as demonstrated in rodents. Here, we performed deep RNA sequencing (RNA-Seq) on laser microdissected CeL neurons collected from brains of phenotyped young rhesus monkeys (n=47). We pooled many individually captured neurons which allows for unbiased estimation of RNA across the transcript. Based on the findings of this study, we neuroanatomically characterized the expression and distribution of a leading AT-related candidate (*PRKCD*) within the extended amygdala in another set of monkeys. This in-depth investigation allows for an understanding of individual differences in CeL neuronal gene expression in relation to individual differences in AT and its components, and points to novel molecular targets for the treatment of anxiety disorders and other stress-related psychopathology.

## Results

### AT phenotyping and CeL neuronal collection

In this study, laser capture microdissection (LCM) was combined with RNA-Seq to identify CeL neuronal gene expression as it relates to individual differences in the AT phenotype and its components (Figure 1A). AT is a composite score of threat-related behavioral and pituitary-adrenal activation occurring during the potentially threatening no eye contact condition (NEC) of the human intruder paradigm (Fox et al., 2010; Fox et al., 2008; Kalin and Shelton, 1989; Kalin et al., 2005.). During NEC, monkeys respond by inhibiting their behavior, as characterized by increases in freezing duration and decreases in coo vocalizations. These responses are accompanied by increases in cortisol, a reflection of hypothalamic-pituitary axis activation. AT is calculated within a sample of animals as a standardized composite of NEC-induced increases in freezing duration, decreases in cooing frequency, and increases in plasma cortisol concentration (see methods).

**Figure 1.**
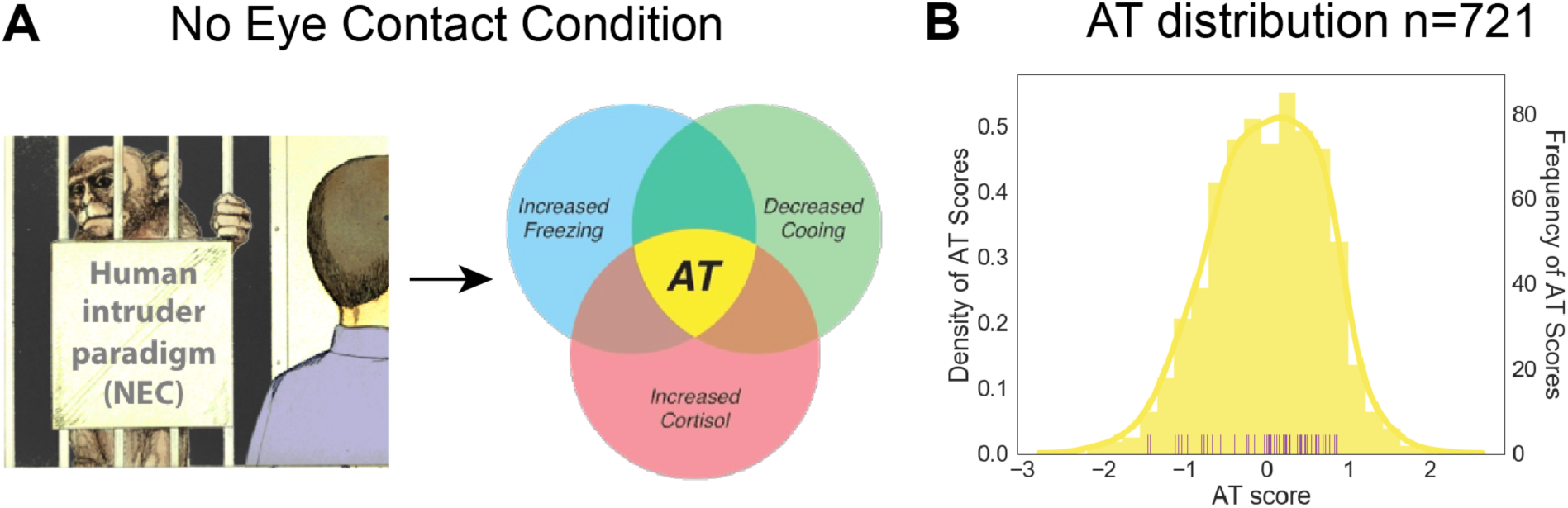
AT phenotype and distribution. (A) Human Intruder No-Eye Contact condition elicits behavioral and physiological responses associated with uncertain threat such as increased freezing, decreased cooing, and increased cortisol, which together comprise the AT-phenotype. (B) Histogram of AT across a larger superset of animals (n=721) in yellow. The purple rugplot below demonstrates the AT scores of the animals in this study as they were calculated within this group of 47 animals.

To determine whether the AT scores of the 47 animals were representative of the AT scores of a much larger population (n=721) from which the animals in this study were a sub-sample, AT was calculated in two ways. First, AT was determined in relation to the 47 animals used in this study and this was compared to the AT scores of these same animals when calculated with the large group of 721 animals. When examining the AT scores of the animals used in this study as calculated in relation to each other, we found that these AT scores had a similar distribution as the AT scores across the whole 721 animals. AT as calculated within the current sample (n=47), spanned approximately +/- 1 std from the mean of the larger cohort (Figure 1B). We computed the ranks of the AT scores for the 47 animals when they were calculated the two different ways and found that the spearman correlation between the ranks was rho=0.7 (p<0.0001). Additionally, a Wilcoxon signed-rank test demonstrated that the ranks did not significantly differ (Z=527, p=0.69). This demonstrates that the AT scores as calculated within the 47 animals reflects the range of individual differences in the larger population.

Previous work from our laboratory demonstrated that metabolism within the Ce is significantly associated with AT (Fox et al., 2015a; Oler et al., 2010). High levels of the serotonin transporter demarcate the CeL so that by imaging the serotonin transporter with positron-emission-tomography, we were able to identify the peak correlation to be within the CeL (Figure 2A; (Shackman et al., 2013). Additionally, neurotoxic lesions of the rhesus Ce result in decreased expression of AT’s components (Kalin et al., 2004; Oler, 2016). These findings in primates, along with rodent studies, formed the basis of our focus on CeL neurons. In the present study, the CeL was identified using acetylcholinesterase staining. In adjacent sections used for LCM, CeL neurons were identified with a neuron-specific rapid staining protocol. This LCM approach was advantageous in specifically identifying the CeL, which changes shape and position across the A-P extent of the amygdala. These CeL neurons were individually captured and pooled, and the RNA was extracted and sequenced (Figure 2B). Across all animals, on average, 90% of the cells that were sequenced were confirmed to have been collected from the CeL, as reflected in our average bregma estimates and CeL neuron accuracy measure for each animal (Supplementary Table 1, Supplementary Figure 1B). Neuronal enrichment was also confirmed, as we found greater expression of neuron-specific genes compared to glia-specific genes (t=36.4, p<0.0001; Supplementary Figure 1C).

**Figure 2.**
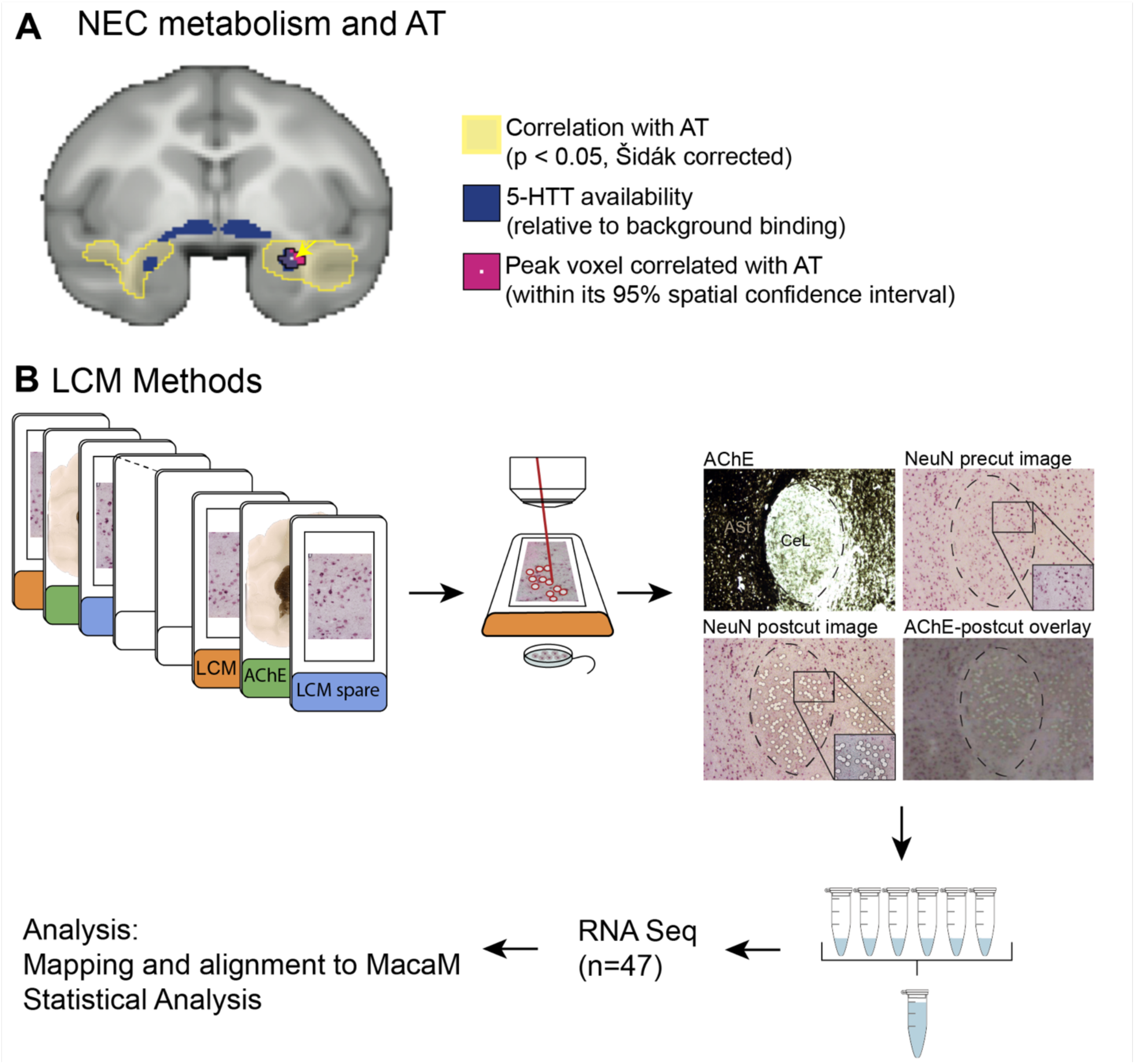
AT-associated PET metabolism in the Ce as a rationale for laser capture microdissection and sequencing of CeL neurons. (A) Using data from our large sample of AT-phenotyped and brain imaged animals, we previously demonstrated that individual differences in NEC-associated PET metabolism are related to individual differences in AT (yellow, (Oler et al., 2010). We identified the peak AT voxels (pink) to be in the Ce region by demonstrating an overlap with serotonin transporter binding (blue), which, relative to surrounding regions, is elevated in the CeL. (B) Brain slabs containing the amygdala were identified and the A-P location was determined before sectioning. Tissue was sectioned and mounted on LCM slides (orange and blue labels). Adjacent slides were stained with AChE (green labels) to determine the location of the CeL. LCM slides were stained with an abbreviated NeuN protocol and CeL neurons were captured into the lid of a microfuge tube. Post-cut overlays were made for every slide to confirm capture location. RNA from 500-600 CeL neurons was pooled and used for RNA-Seq (n=47). Reads were processed and aligned to MacaM (Zimin et al., 2014).

### AT as a predictor of CeL neuronal gene expression as compared to its individual components

The importance of the AT construct is that it combines behavioral and physiological responses that are associated with threat (Fox et al., 2008). The co-occurrence of these responses has been implicated in the expression of anxiety disorders and other stress-related psychopathologies (Chronis-Tuscano et al., 2009; Davidson and Rickman, 1999; Essex et al., 2010; Fox and Kalin, 2014; Kagan et al., 1987). Additionally, at the level of the amygdala, the CeL functions to coordinate the co-expression of these behavioral and physiological responses aimed at promoting survival (Veening et al., 1984; Viviani et al., 2011). Previous work from our laboratory assessing regional brain metabolism in relation to AT and its components demonstrated that AT accounts for a greater amount of variance in CeL metabolism than any one of its components alone (Shackman et al., 2013). Based on this, we explored the hypothesis that AT would be a more robust, transcriptome-wide, predictor of gene expression compared to each of its components.

This LCM technique allowed us to acquire and sequence sufficient amounts of RNA such that a multiple regression, individual differences approach could be used to investigate the relationship between gene expression and each predictor of interest. Transcripts were identified based on whole exon gene expression data that were quantile normalized and log2 transformed. We first tested the extent to which AT, as the predictor in our model, performs above chance. A simulated null distribution was constructed by shuffling AT and determining how many genes were significantly associated with AT and repeating this process 10,000 times (Figure 3A). Results demonstrated that AT performed significantly better than chance (empirical p = 0.04, Figure 3B). We also tested whether each of the individual components of AT predicted significantly more genes above chance (Figure 3A) and found that the number of genes observed to be associated with each component was not significantly greater than the mean of the simulated distribution for each component (empirical p-values for freezing: p=0.3, cooing: p=0.24, cortisol: p=0.12; Figure 3B). Additionally, we found that AT significantly predicted more genes above chance compared to each of AT’s components (t-test of AT compared to: freezing t=113.5, p<0.001, cooing t=106.9, p<0.001, and cortisol t=496, p<0.001; Figure 3C). We also examined the overlap among AT-related genes and those associated with freezing, cooing, and cortisol (Figure 3D). Five hundred and fifty-five genes were significantly associated with AT with 383 genes also being significantly associated with at least one of AT’s components, leaving 172 genes that were significantly correlated with AT and not one of its components. Taken together, these analyses indicate that the AT phenotype captures additional variance in gene expression beyond that accounted for by its individual components. This suggests that there are some CeL genes that are associated with the AT phenotype without being related to a specific way in which AT is expressed.

**Figure 3.**
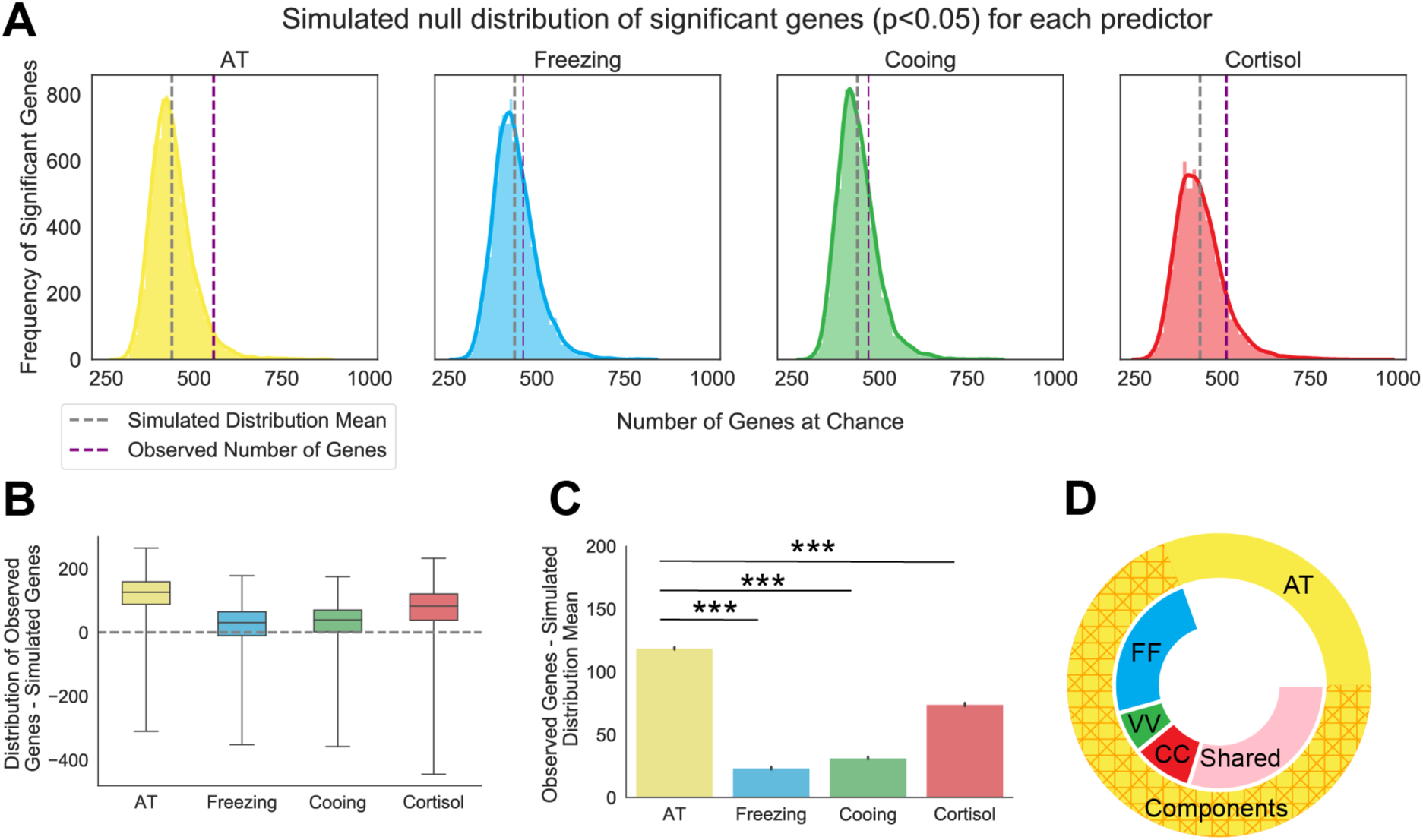
AT predicts a significant number of genes above chance and outperforms each of the AT components. (A) Simulated null distribution (as described in the methods section RNA Sequencing Analysis and Statistics) for each predictor of interest at a nominal p-value of p<0.05. Purple dotted lines indicate the observed number of genes associated with each predictor’s real values. Grey dotted lines indicate the mean number of genes of the simulated distribution. Solid colored outline of the distribution represents the density of significant genes as determined by a kernel density estimation. (B) Boxplots for each predictor depicting the distribution and mean of the differences between the real observation and each simulated value. Empirical p-values were calculated for each predictor, AT: p=0.04, freezing: p=0.3, cooing: p=0.24, cortisol: p=0.12 (C) Barplot demonstrating that AT predicts significantly more genes above the simulated distribution mean than those predicted by each of the AT components alone: freezing (t=113.5, p<0.001), cooing (t=106.9, p<0.001), or cortisol (t=496, p<0.001). Error bars are displayed as SEM. P-values are Šidák corrected for multiple comparisons. AT component values were transformed and residualized as described in the Methods: Anxious Temperament phenotyping (D) Donut plot depicting the number of overlapping genes between individual AT components and AT. Outside circle represents all 555 AT-related genes (p<0.05) and is separated into the 393 genes that overlap with AT components (hashed orange) and the 172 genes that are unique to AT (yellow). Inner circle represents genes that are related to AT and is broken up by genes that are also unique to one AT component (FF: freezing in blue, VV: cooing in green, CC: cortisol in red) or that are shared by at least two components (shared in pink).

To expand our analyses of the transcriptomic architecture of AT as it relates to its components, we performed four separate regression analyses for each gene’s expression level in relation to our phenotypic measures: AT, freezing, cooing, and cortisol. We then asked whether the t-values from each of these regressions between genes and phenotype were correlated across components and AT. Results demonstrated that significant correlations were found between the t-values characterizing the relation between expression levels and AT with the t-values for freezing (Supplementary Figure 3A, rho=0.88, p<0.001), cooing (Supplementary Figure 3B, rho=-0.72, p<0.001), and cortisol (Supplementary Figure 3C, rho=0.33, p<0.001). When comparing gene relations among components, we also found associations between genes related to freezing with those related to cooing (Supplementary Figure 3D, rho=-0.76, p<0.001) as well as between genes associated with cortisol and cooing (rho=0.31, p<0.001, Supplementary Figure 3E). We did not observe this relationship between genes associated with freezing and cortisol (Supplementary Figure 3F, rho=-0.02, p=0.27). These data support the hypothesis that AT-related genes include component-general genes such as those that are associated with multiple components and AT rather than a single component, as well as component-specific genes, e.g. those specifically related to freezing.

### CeL neuronal transcripts associated with AT

Multiple regression analyses demonstrated 555 AT-related transcripts significant at a p-value of p<0.05 with 231 negative relationships and 324 positive relationships (two-tailed uncorrected; top 100 genes in Figure 4A; Supplementary Table 3). Of these 555, the relation between AT and 528 transcripts passed permutation testing in which 10,000 correlations were run for each gene with randomly shuffled AT scores. Eighty percent of the top 100 transcripts that passed permutation testing were confirmed as significantly associated with AT in a separate analysis using the same model implemented in DESeq2. Based on gene ontology (GO) families, AT-related genes were characterized as being predominately located within the cell body and fell into a number of different gene families that included molecular binding and developmental processes (Supplementary Table 4). GO enrichment analysis also revealed a variety of biological processes associated with AT (Supplementary Table 5). For example, a number of transcripts were identified that fell into the GO categories of neuron projection (GO:0043005), cell projection regulation (GO:0031344), and G-protein receptor activity (GO:0031344; Figure 4B).

**Figure 4.**
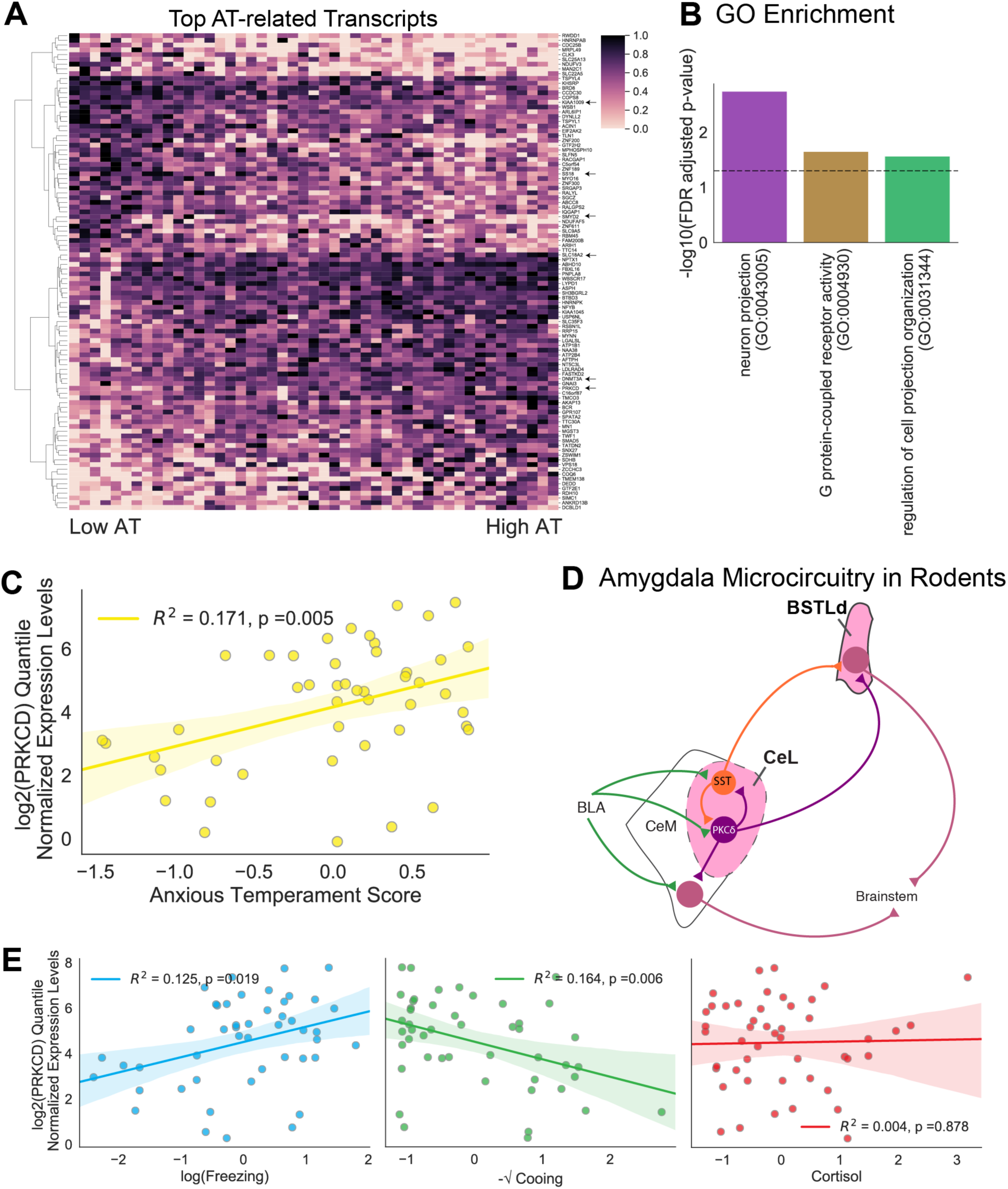
RNA-Seq of LCM CeL neurons revealed AT-related genes. (A) Heatmap of the top 100 transcripts related to AT (p<0.05, passing permutation testing, OLS regression). Gene expression data are presented as quantile normalized min-max scaled values. Arrows point to genes that are discussed in the text. (B) Most significant GO enrichment groups for cellular component (purple), molecular function (yellow) and biological process (green). The FDR-corrected p-value is depicted by the black dashed line. (C) Correlation between PRKCD mRNA expression levels and AT (R^2^=0.171, p=0.005). Shaded areas represent the SEM. (D) Simplified diagram of the microcircuit within the rodent amygdala and extended amygdala. PKCδ expressing neurons are labeled in purple. SST expressing neurons are labeled in orange. Previous work demonstrates that SST and PKCδ expressing neurons both receive information from the basolateral amygdala complex (BLA) and contribute to an inhibitory microcircuit within the CeL (Haubensak et al., 2010; Li et al., 2013; Ye and Veinante, 2019). Both cell types also project to the laterodorsal BST (BSTLd) while PKCδ expressing neurons but not SST neurons project to the CeM (Haubensak et al., 2010; Li et al., 2013; Ye and Veinante, 2019). (E) Correlations between *PRKCD* mRNA expression levels and the individual components of AT (freezing; R^2^=0.125, p=0.019, cooing; R^2^=0.164, p=0.006, cortisol; R^2^=0.004, p=0.875; OLS regression). Freezing, cooing, and cortisol values were standardized, transformed, and residualized as described in the methods. *PRKCD* mRNA expression levels are presented as quantile normalized log2 transformed values residualized for age at ToD and CeL neuron accuracy.

More specifically, we identified new transcripts as well as transcripts of interest based on previous literature (Figure 4A). Among the novel transcripts, we found several that are related to epigenetic mechanisms such as the SS18 subunit of BAF chromatin remodeling complex (*SS18;* t = −4.79, p= 0.00002), histone methyltransferase SET and SMYND domain containing 2 *(SMYD2;* t = −3.53, p= 0.001), and the DNA methyltransferase 3 (*DNMTA3;* t = 3.57, p= 0.0009). This finding is intriguing in light of our previous work demonstrating a lack of Ce metabolism co-heritability with AT and points to genes within the CeL that could mediate the non-heritable factors (e.g. environment) of AT-related Ce function. Another novel transcript was Centrosomal Protein 162 (gene name: *KIAA1009* or *CEP162*; t= −4.39, p= 0.00007). This is an interesting candidate as it plays a role in primary cilia function in adult cells (Wang et al., 2013) and recent preclinical and *in vitro*-derived neuronal studies have found alterations in primary cilia to be linked to schizophrenia (Munoz-Estrada et al., 2018) and cognitive function (Amador-Arjona et al., 2011; Berbari et al., 2014; Wang et al., 2011). Transcripts of interest based on previous literature included: neuropeptide Y receptor 2 (*NPY2R*, t=2.39, p=0.0213), solute carrier family 18 member A2 (gene name: *SLC18A2* or *VMAT-2;* t = 3.79, p=0.00046) and protein kinase C-type delta (gene name: *PRKCD*, protein name: PKCδ; t= 2.93, p=0.0052; Figure 4A, C). *NPY2R* has been previously associated with the amygdala and anxiety-related behaviors (Carvajal et al., 2006; Tschenett et al., 2003) and *VMAT-2* plays an important role in monoaminergic neurotransmission (Frank, 2009, 2010; Guay, 2010; Taylor et al., 2009).

*PRKCD* is a particularly interesting candidate because it is a marker for a CeL neuron population that is involved in threat-processing and conditioned fear learning in rodents (Figure 4D; (Cui et al., 2017; Haubensak et al., 2010; Yu et al., 2016). For example, CeL *PRKCD* expressing neurons decrease their firing in response to a conditioned stimulus, thereby disinhibiting CeM neurons, which results in increased freezing behavior (Haubensak et al., 2010). Within the cell, PKCδ serves as a node by which many secondary messenger pathways intersect and mediates numerous processes (Newton, 2010) including neurotrophic signaling which has been linked to AT and neuropsychiatric disorders (Duman and Li, 2012; Fox et al., 2012; Fox et al., 2019; Turner et al., 2011). In relation to the individual components of AT, *PRKCD* mRNA expression was significantly associated with increased freezing (t=2.44, p= 0.019) and decreased cooing (t=-2.87, p= 0.006) but not threat-related cortisol (t=0.15, p= 0.875; Figure 4E), suggesting it may be more strongly associated with the behavioral inhibition-related components of AT.

### Characterizing PKCδ and SST neurons in the nonhuman primate CeL

While PKCδ has been extensively studied in the rodent CeL (Amano et al., 2012; Cui et al., 2017; Haubensak et al., 2010; Ye and Veinante, 2019; Yu et al., 2016), little is known about its expression in the nonhuman primate amygdala. Rodent studies demonstrate that other CeL neurons exist (Cassel and Gray, 1989; McCullough et al., 2018; Price et al., 1987; Roberts et al., 1982) and recent mouse studies reveal that *SST* expressing GABAergic neurons interact with and modulate the function of *PRKCD* expressing neurons (Fadok et al., 2017; Kim et al., 2017; Li et al., 2013). Because these cell types have not been well characterized in the nonhuman primate, we used stereological cell counting techniques to map the distribution of PKCδ and SST expressing neurons in the rhesus CeL. To further understand the extent to which the mouse CeL microcircuitry studies are translatable to primates, we also performed parallel studies in the mouse CeL (Figure 5). In the nonhuman primate CeL, PKCδ expressing neurons accounted for 59% of the estimated number of neurons and SST expressing neurons accounted for only 6% (Figure 5D). In the mouse CeL, PKCδ expressing neurons constituted 43% of the total estimated number of neurons and SST expressing neurons accounted for 20% (Figure 5D). While the proportion of PKCδ expressing cells to neurons did not significantly differ between species (t=-1.06, p=0.34; Figure 5E), the proportion of SST expressing cells to total neurons were notably decreased in the monkey compared to the mouse (t=-3.6, p=0.02; Figure 5E). In nonhuman primates, 4% of neurons expressed both SST and PKCδ while in mice, this population was virtually non-existent (Figure 5D).

**Figure 5.**
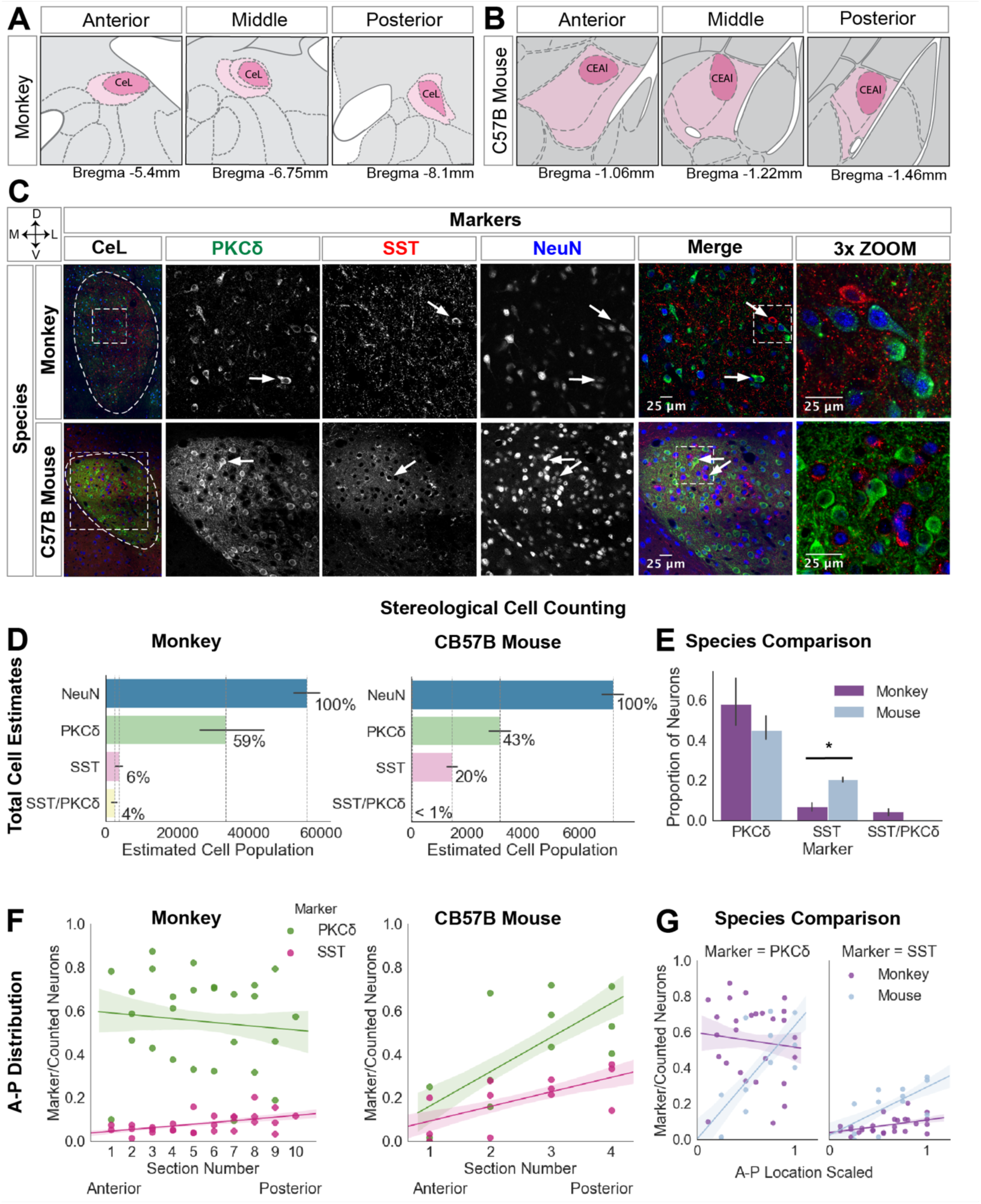
PKCδ expressing neurons in the monkey compared to the mouse CeL. (A) CeL atlas slices depicting the A-P extent in the rhesus monkey (Paxinos et al., 2009); (B) and in the mouse (Allen Brain Atlas). (C) Representative confocal images of the CeL in both species; white arrows point to PKCδ and SST neurons. Images were adjusted using the Fiji despeckle filter (Schindelin et al., 2012) for removing salt and pepper noise. (D) Stereological cell estimates for monkey (n=3) and mouse (n=3). (E) Species comparison of PKCδ, SST, and PKCδ/SST estimates are presented as a proportion of the total number of neurons (PKCδ: t=-1.06, p=0.34, SST: t=3.6, p=0.02; t-test), error bars are SEM. (F) A-P distribution of PKCδ and SST expressing neurons in monkey and mouse (monkey: PKCδ p=0.7, t=-0.39 and SST p=0.012, t=2.7, mouse: PKCδ p=0.01, t=3.1, SST p=0.02, t=2.6; OLS regression). (G) Species comparison of the A-P distribution of each cell type (A-P location X species interaction for PKCδ (t=2.6, p=0.01) and A-P location X species interaction for SST (t=2.7, p=0.01); OLS regression). To compare A-P distribution between species, A-P location was min-max scaled with 0 indicating more anterior slices and 1 indicating more posterior slices.

Previous studies in mice demonstrate that cell type markers are differentially distributed across the anterior-posterior (A-P) extent, perhaps suggesting functional differences between the anterior and posterior Ce (Amaral et al., 1989; Han et al., 2017; Haubensak et al., 2010). Consistent with this, we found that both PKCδ (t=3.1, p=0.01) and SST (t=2.6, p=0.02) expressing neurons were significantly more concentrated in the posterior mouse CeL (Figure 5F). In the monkey we found that the SST expressing somata were more concentrated in the posterior region of the CeL (t=2.7, p=0.012; Figure 5F), replicating our previous observation in the nonhuman primate (Kovner et al., 2019). However, deviating from the mouse, nonhuman primate PKCδ expressing neurons were not differentially distributed across the A-P extent of the CeL (t=-0.39, p=0.7). A two-way interaction between A-P location and species was tested separately for PKCδ neurons and SST neurons. As can be seen in Figure 5G, the slopes of the lines for A-P location for both PKCδ neurons (t=2.6, p=0.01) and SST neurons (t=2.7, p=0.01) differed between species.

In contrast to the relatively small number of SST expressing neurons in the CeL, and consistent with previous work, we found dense SST neuropil throughout the nonhuman primate CeL (Amaral et al., 1989; Cassell et al., 1986; Kovner et al., 2019; Martin et al., 1991). Numerous SST varicosities were present, and we were able to identify these SST varicosities in close apposition to the primary dendrite and cell soma of some large CeL neurons, a number of which expressed PKCδ (Figure 6D). This finding suggests that SST input modulates CeL PKCδ expressing neurons in nonhuman primates. Compared to the relatively limited distribution profile described in the mouse (Haubensak et al., 2010), we observed that PKCδ expression in the monkey was widely distributed across the brain (Supplementary Figure 4B). This finding was further confirmed with human and nonhuman primate microarray data from the Allen Brain Atlas (Supplementary Figure 4A).

**Figure 6.**
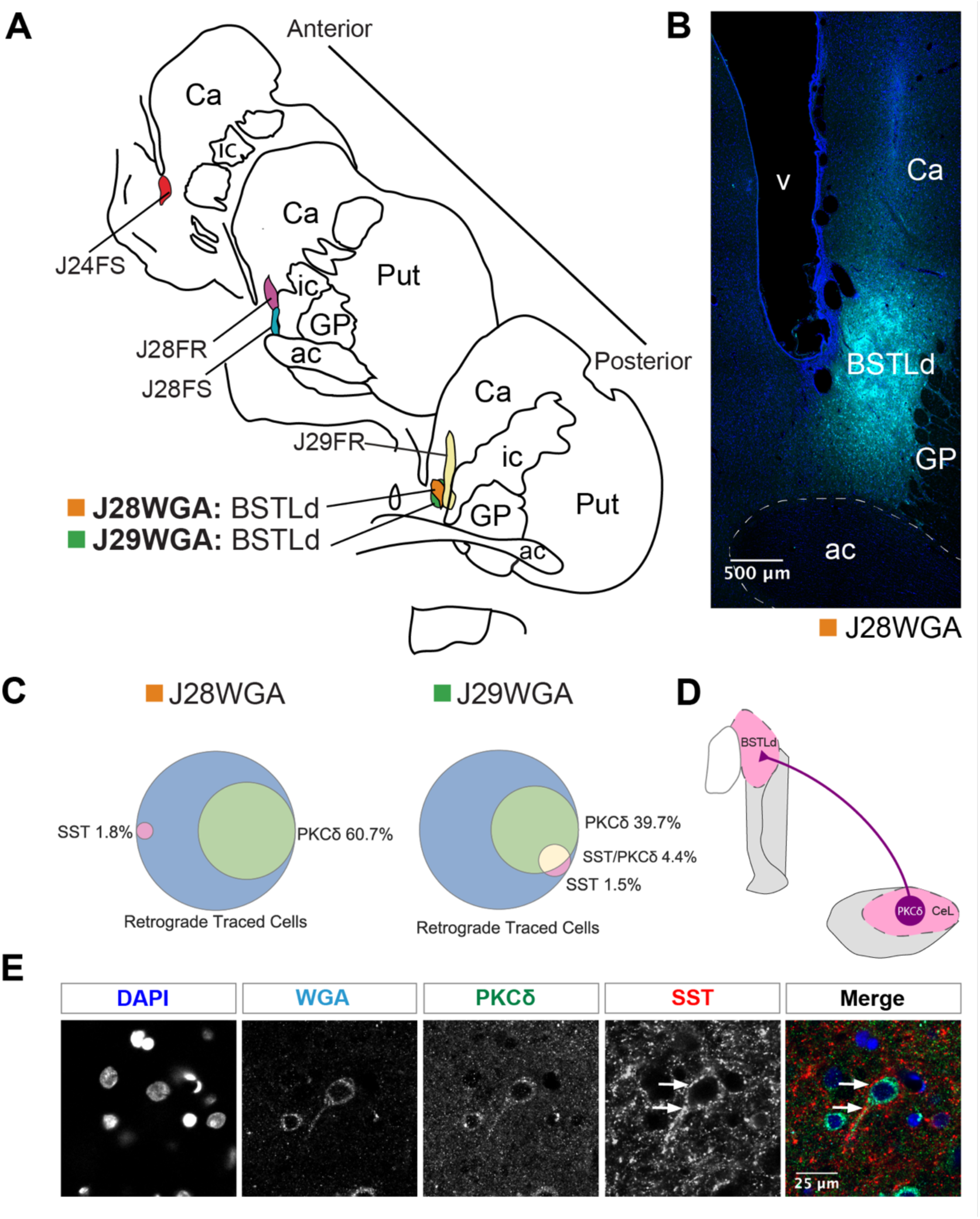
A subset of PKCδ expressing neurons project to the BSTLd in the monkey. (A) Hand drawn slices depicting the localization of retrograde tracer into different regions of the BST in monkey. Two replicates, J29WGA and J28WGA, are localized to the same part of the BSTLd. (B) Representative confocal image of the BSTLd injection site. DAPI staining is in blue. WGA-tracer staining is in cyan. (C) A Venn diagram for each BSTLd replicate, J29WGA and J28WGA, illustrating the percent overlap between the WGA-tracer and PKCδ, SST, or both. (D) Simplified diagram of our results demonstrating that CeL PKCδ expressing neurons project to the BSTLd in nonhuman primates. (E) Representative confocal image of a BSTLd-projecting neuron that expresses PKCδ. This image was adjusted using the Fiji despeckle filter (Schindelin et al., 2012) for removing salt and pepper noise. White arrows point to the immense SST innervation received along this neuron’s primary dendrite and soma.

### A subset of CeL PKCδ neurons project to the BSTLd in the nonhuman primate

In rodents, in addition to constituting a CeL threat-related microcircuit, *PRKCD* and *SST* expressing neurons project to other parts of the extended amygdala (Ahrens et al., 2018; Cai et al., 2014; Ye and Veinante, 2019). For example, *PRKCD* expressing neurons have short range projections to the CeM and both *PRKCD* and *SST* expressing neurons project to the laterodorsal BST (BSTLd; which corresponds to the oval nucleus of the BST in mice; Figure 4D; (Ahrens et al., 2018; Cai et al., 2014; Haubensak et al., 2010; Ye and Veinante, 2019). This suggests that CeL PKCδ and SST expressing neurons may serve to coordinate CeL and BSTLd in mediating threat-related behaviors. Because there are differences in intrinsic extended amygdala connectivity between nonhuman primates and rodents (Oler et al., 2017), it is important to characterize the extent to which CeL PKCδ and SST expressing neurons project to the BSTLd in nonhuman primates. In six cases involving nonhuman primates, retrograde tracers were injected into different subregions of the BST and in two of these cases the injections were centered in the BSTLd (J29WGA and J28WGA; Figure 6A-B). Tissue was co-labeled for the retrograde tracer, DAPI, PKCδ, and SST (Supplementary Table 2). As seen in our previous work with other tissue from these animals (Oler et al., 2017), the cases with injections directly into the BSTLd (J29WGA and J28WGA; Figure 6A-B) demonstrated substantially more CeL retrograde labeled cells (Table 1). Because of our specific interest in BSTLd projecting CeL neurons, we focused on these two cases. Across these cases, CeL retrograde-labeled cells expressing PKCδ ranged from 40-60% (Figure 6C). In contrast, very few retrograde-labeled cells exclusively expressed SST or co-expressed SST and PKCδ (Figure 6C). Adding to our previous observation of SST varicosities surrounding CeL somata and primary dendrites, we also observed that this was present in some CeL to BSTLd projecting neurons, a subset of which expressed PKCδ (Figure 6D). These data suggest that of the two populations investigated here, it is primarily PKCδ neurons that project to the BSTLd, and that SST neurons are well-poised to inhibit this projection.

**Table 1.**
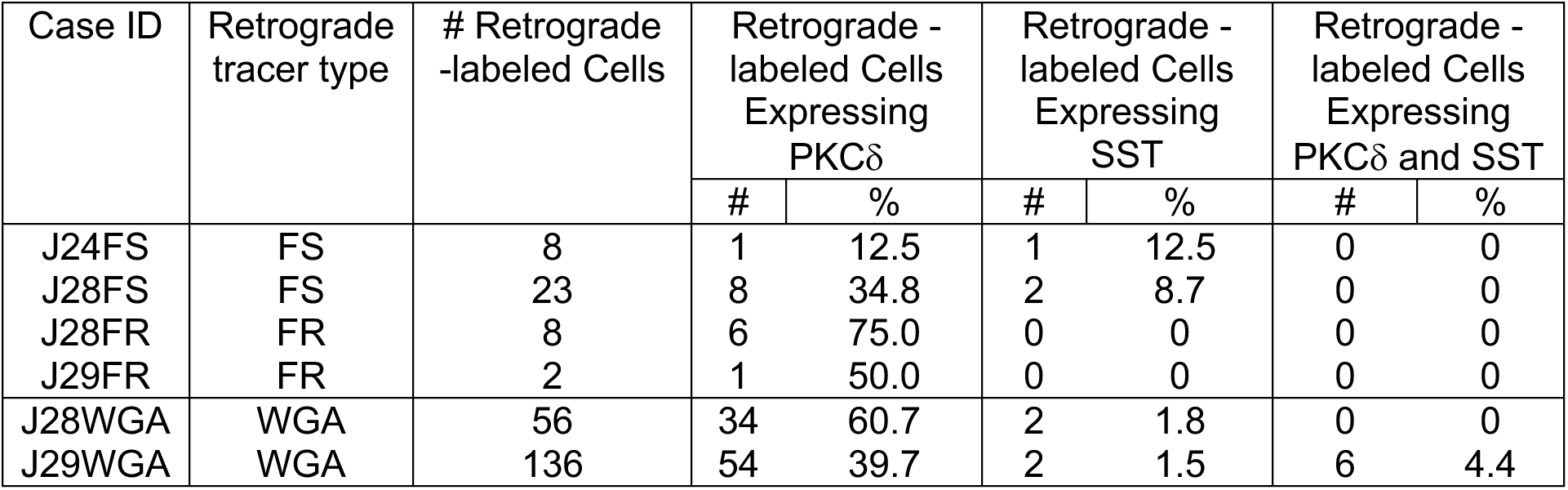
Number and percentage of retrograde tracer-labeled cells expressing markers of interest for each case of retrograde injection.

## Discussion

In our previous work we demonstrated the importance of the primate Ce in mediating AT, the early-life risk phenotype underlying the development of anxiety and depression. Here, we build on these studies by characterizing alterations in gene expression in neurons from the CeL, a subdivision of the Ce that is critical for gating the flow of anxiety-related information within the amygdala (Fadok et al., 2018). By performing RNA-Seq on mRNA extracted from LCM captured CeL neurons from 47 animals fully phenotyped for AT, we identified numerous transcripts that are associated with AT and that are components of several biological and molecular processes. *PRKCD* is one of the more interesting AT-related transcripts since rodent studies demonstrate a role for CeL PKCδ expressing neurons in modulating threat-related freezing behavior, a primary component of AT (Haubensak et al., 2010; Yu et al., 2017).

Our whole-transcriptome analysis demonstrated that AT is able to predict gene expression above chance and does this more robustly than each AT component individually. This is consistent with previous work from our laboratory demonstrating that combining AT components into a single composite measure accounts for more variance in brain metabolism as compared to each component alone (Shackman et al., 2013). Overall, the RNA-Seq data demonstrated that 6% of the transcripts, or 555 out of the 8747 transcripts investigated, were associated with individual differences in AT. Consistent with our earlier work, the ontology analysis of our neuronal CeL sample overlapped with previously identified AT-related pathways (Fox et al., 2012; Fox et al., 2019).

With our large multigenerational pedigree, we previously examined the co-heritability of AT-related brain metabolism with the AT phenotype. Results demonstrated significant co-heritability of AT with threat-related metabolism in the BST, posterior OFC, and periaqueductal gray, but not in the Ce. This suggests that Ce metabolism as it relates to AT may be particularly influenced by non-heritable factors. In this regard, it is interesting that our CeL neuronal data point to several transcripts involved in epigenetic mechanisms (e.g. *SS18*, *SMYD2*, and *DNMTA3*). These genes have been implicated in DNA methylation, histone methylation, and chromatin remodeling pathways (Li et al., 2018; Lubieniecka et al., 2008; Tang et al., 2010) and could play a role in linking environmental experience to CeL neuronal function and plasticity.

*KIAA1009* (*CEP162*), another interesting candidate gene that was strongly associated with AT, codes for a protein that is located at the base of primary cilia (Wang et al., 2013). Although this gene has not been studied in the context of neuropsychiatric disorders, alterations in primary cilia structure have been implicated in reduced adult neurogenesis (Amador-Arjona et al., 2011) and poor novel object learning (Wang et al., 2011), and fewer primary cilia are observed in neurons from schizophrenic patients that were derived *in vitro* (Munoz-Estrada et al., 2018). In the context of our data, these studies provide a rationale for additional work examining the role of *KIAA1009* in mediating altered cilia structure and function as it relates to stress-related psychopathology (Pruski and Lang, 2019). Increased *NPY2R* was also associated with AT (t = 1.93, p=0.0256) and is consistent with previous studies demonstrating the importance of the *NPY* system in mediating fear and anxiety responses (Roseboom et al., 2014). For example, *NPY2R* agonist administration into the amygdala increase anxiety-like responses (Sajdyk et al., 2002) and that global or Ce-specific *NPY2R* knock out (KO) mice express reduced anxiety-like behaviors (Carvajal et al., 2006; Tasan et al., 2010; Tschenett et al., 2003). Based on our prior work, we expected to observe an association between AT and neurotrophic kinase receptor type 3 (*NTRK3*) gene expression (Fox et al., 2012; Fox et al., 2019). In our recent study, we identified an association between individual differences in *NTRK3* exon 3 expression with AT, indicating a role for the truncated isoform of *NTRK3* in AT (Fox et al., 2019). Unfortunately, the amount of input RNA from our current LCM study did not allow us to examine *NTRK3* expression at this gene feature level, which could explain this discrepancy.

An exciting finding from the current study is the observation that greater *PRKCD* mRNA expression is associated with increased AT. While it is likely that these changes would be reflected in PKCδ protein levels, we acknowledge the possibility of a dissociation between mRNA and protein. Because of the methods used to collect LCM neurons, we were unable to assess protein levels of PKCδ in the animals used for RNA-Seq. Studies in the mouse demonstrate that CeL *PRKCD* expressing neurons are a component of a microcircuit that gates CeL output, modulating the expression of freezing and other threat-related behaviors (Ciocchi et al., 2010; Fadok et al., 2017; Haubensak et al., 2010; Li et al., 2013; Yu et al., 2016). Because little is known in primates about the proportion and distribution of CeL neurons that express PKCδ protein, in three other animals, we systematically characterized CeL PKCδ expressing neurons. Results demonstrated that PKCδ expressing neurons accounted for 59% of total CeL neurons and that these neurons were evenly distributed across the AP extent of the CeL. Because we found that a portion of CeL neurons expressed PKCδ, the possibility exists that this population could be greater in number or that PKCδ has greater functionality in high AT animals, thereby accounting for the association between *PRKCD* mRNA and AT.

Studies in rodents and nonhuman primates demonstrate that the CeL sends major projections to the BSTLd and these extended amygdala components are directly linked to the expression of fear and anxiety-related responses (Ahrens et al., 2018; Oler et al., 2017; Petrovich and Swanson, 1997; Pomrenze et al., 2019; Ye and Veinante, 2019). Therefore, we were particularly interested in the extent to which CeL PKCδ neurons project to the BSTLd. Not only did we find that PKCδ neurons project to the BSTLd, they constituted approximately half of the retrograde-labeled neurons that were identified. Based on this observation, the association between *PRKCD* mRNA expression in the CeL and AT, and the fact that BST is part of the AT circuit, it is conceivable that these neurons are critical in mediating the flow of information relevant to the expression of AT. Future studies will be necessary to test this hypothesis. It is also noteworthy that approximately half of the retrograde-labeled neurons did not express PKCδ and suggests that other CeL neuron populations project to the BSTLd, such as those expressing corticotropin-releasing factor (Gray and Magnuson, 1992; Moga and Gray, 1985; Pomrenze et al., 2019).

In general, our findings are consistent with the elegant studies performed in rodents pointing to the importance of CeL PKCδ expressing cells in regulating threat responding. To date, these studies have examined the role of PKCδ expressing cells, however the actual role of PKCδ protein in relation to threat-processes has not yet been investigated. Other studies demonstrate PKCδ’s involvement in the PLC/PIP2/DAG pathway, a commonly used secondary messenger system of neurotrophic signaling (Rankin et al., 2008), chemokine signaling, and membrane steroid signaling (Greene et al., 2004; Kanehisa and Goto, 2000; Kanehisa and Sato, 2019). While the molecular function of PKCδ in relation to fear and anxiety is still unknown, our work confirms that targeting CeL PKCδ has high translational potential as a therapeutic target. Studies manipulating CeL PKCδ expression will be useful for understanding PKCδ’s role in mediating behaviors associated with threat-processing.

*SST* expressing neurons also exist in the CeL and rodent studies demonstrate an interaction between *SST* expressing neurons and *PRKCD* expressing neurons in modulating CeL output (Fadok et al., 2017; Haubensak et al., 2010; Li et al., 2013). While we did not find *SST* mRNA to be associated with AT, because of its potential modulatory role, we systematically characterized the population of SST expressing neurons in the primate CeL using immunofluorescence staining. By comparing tissue from the monkey and mouse, we found that in the monkey, SST expressing neurons constituted a much smaller population than was observed in the mouse. Consistent with previous work (Amaral et al., 1989; Cassel and Gray, 1989; Martin et al., 1991), we also noted dense CeL SST neuropil in both the nonhuman primate and the mouse. It is noteworthy that in the nonhuman primate CeL, we observed neurons that were surrounded by SST varicosities, some of which expressed PKCδ and projected to the BSTLd. We also observed that SST varicosities appear to innervate the primary dendrite of some CeL neurons. The origin of the SST innervation in the nonhuman primate CeL is not known. However we do know that SST is predominantly expressed in a subset of GABAergic neurons and that there are a limited number of GABAergic regions that send input specifically to the CeL (Oler et al., 2017; Petrovich and Swanson, 1997). Therefore, it is plausible that the CeL SST innervation originates from other parts of the central extended amygdala (e.g. BST or sublenticular extended amygdala) and/or projections from the amygdala intercalated cell masses (Ahrens et al., 2018; Gungor et al., 2015; Pare and Smith, 1993). That said, the small number of CeL SST neurons may also be largely responsible for the dense SST CeL neuropil. This is based on rodent studies demonstrating that CeL SST neurons project to other CeL neurons (Cassell et al., 1986; Gray et al., 1982), including those that express PKCδ (Fadok et al., 2017; Li et al., 2013). To further understand the organization and functional implications of this circuitry in nonhuman primates, future studies should determine the origins of CeL SST input and investigate the effects of SST release on CeL PKCδ neurons in mediating threat-related behavior.

This transcriptome-wide study in nonhuman primate CeL neurons provides a molecular basis for understanding amygdala alterations as they relate to the early-life risk to develop anxiety disorders and related psychopathology. This is the first study to fully characterize gene expression in nonhuman primate CeL neurons and to implicate CeL PKCδ expressing neurons in primate AT. To provide a more in depth understanding of CeL PKCδ expressing neurons in primates, we systematically characterized these neurons and by comparing them to the mouse, we found potentially relevant species differences. Importantly, we demonstrate that some of the CeL PKCδ expressing neurons in the nonhuman primate project to the BSTLd and that some of these neurons are innervated by SST varicosities. These findings present evidence in the primate of a neural circuit that maybe highly relevant to understanding human anxiety and point to specific molecules within this circuit that could serve as potential treatment targets for anxiety disorders.

## Acknowledgements

This work was supported by funding awarded to NHK from the NIMH (R01MH081884 and R01MH046729), and R01MH063291 awarded to JF, and RK from the NIMH (5T32MH018931), grants to the Wisconsin National Primate Center Research Center (P51-OD011106 and P51-RR000167), and the California National Primate Research Center (P51OD011107). Confocal microscopy was performed at the University of Wisconsin-Madison Biochemistry Optical Core, which was established with support from the University of Wisconsin-Madison Department of Biochemistry Endowment. We thank the Neuroscience Training Program at UW-Madison, the personnel of the Harlow Center for Biological Psychology, the HealthEmotions Research Institute, the Waisman Laboratory for Brain Imaging and Behavior, the Wisconsin National Primate Research Center, the Wisconsin Institutes for Medical Research, S. Shelton and H. Van Valkenberg. We thank L. Kordyban, A. Meier, J. Schnabel, A. Elhers for help with LCM, M. Kenwood for assistance with collating and analyzing the population behavioral data, K. Peelman for help with the mouse perfusions.

## Methods

### Experimental Design

Forty-seven animals (average age = 2.27, std = 0.46, 24 males and 23 female) were used for RNA-Seq in this study. To understand the AT levels in these 47 animals in relation to a larger population from which they came, we performed an analysis pooling animal that were phenotyped in our laboratory over the last 12 years. This resulted in a sample of seven hundred and twenty-one preadolescent rhesus monkeys (*Macaca mulatta*), 90 of which were not part of previous publications. These 721 animals were an average age of 1.9 years old (std = 0.74 years) and included 386 males and 335 females. All animals were phenotyped for AT as described below and brains for the subset of 47 animals were collected for LCM and RNA-Seq analysis. In the subset of 47 animals, the males were phenotyped twice and the females once. Phenotyping data that was closest to the time of death was used for the RNA-Seq analysis. Two C57BL/6J mice, two KO; B6;129X1-*Prkcd^tm1Msg^*/J mice, and two additional rhesus monkey brains were used to characterize antibodies. Triple-labeling immunofluorescence and stereology was performed on three C57BL/6J mice (WT, age=p21) and three rhesus monkey brains (average age= 3.1yrs). Triple-labeling immunofluorescence for the retrograde traced issue was performed on six monkeys (*Macaca fascicularis* and *Macaca nemestrina*; average age= 2-3 years).

### Animal Housing

Animal housing and experimental procedures were in accordance with institutional guidelines. All procedures were approved by the Committee on the Ethics of Animal Research of the University of Wisconsin and the Committee on Animal Research at the University of Rochester. Veterinary care, supported by clinical laboratories, was available at the both the Wisconsin National Primate Center and the Harlow Center for Biological Psychology, as well as the University of Rochester 24 hours a day, seven days a week. Monkeys were mother-reared, and pair-housed in a room containing multiple cages, to facilitate social housing. Monkeys experienced a 12-hour light/dark cycle, received food 1-2 times per day and water was available *ad libitum*. Monkeys experienced enrichment activities such as toys, foraging devices, tactile devices, audio, television and snack foods such as fruit at least once per day. Mice were order from The Jackson Laboratory (Bar Harbor, ME) and were group housed at the Wisconsin Psychiatric Institute and Clinics for at least 7 days before experimentation. Mice experienced a 12-hour light/dark cycle and were given *ad libitum* access to food and water. Any discomfort, distress, pain, and injury were minimized by the appropriate use of anesthetic and analgesic drugs under the direction and supervision of the veterinary staff.

### Anxious Temperament phenotyping

Anxious temperament (AT) phenotyping was performed between 8:00am and 11:00am to control for time of day. To assess individual differences in AT, we used the mildly threatening no eye contact condition (NEC) of the human intruder paradigm (Fox et al., 2010; Fox et al., 2008; Kalin and Shelton, 1989; Kalin et al., 2005.) During this 30 minute condition, a human intruder presents their profile to the monkey ensuring to not make eye contact (Fox et al., 2008; Kalin and Shelton, 1989). This condition is unique in that it elicits behaviors that reflect uncertainty in relation to potential threat. High levels of AT are characterized by increased plasma cortisol, increased freezing, and decreased cooing occurring during the NEC condition (Kalin and Shelton, 1989). Freezing is defined as a 3 second period characterized by lack of body movement that is accompanied by a tense body posture and sometimes slow movements of the head. Coo vocalizations are vocalizations made by rounding and pursing the lips characterized by an increase, then decrease in frequency and intensity. The duration of freezing was assessed for six consecutive 5 minute blocks. The average freezing duration across the blocks was log transformed to ensure a normal distribution. Cooing frequency was similarly averaged across the blocks and square root transformed. Blood samples for cortisol assessment were collected by femoral venipuncture at the end of the 30-minute NEC exposure. FDG PET imaging was performed at the end of the NEC period but is not included in this paper. Because cortisol is affected by time of day, cortisol levels were residualized for the time of day of sample collection. Freezing, cooing and cortisol were also residualized for age. For the 47 animals, AT scores were calculated by taking the average of their combined residualized and standardized freezing, cooing, and cortisol values. When examining the AT scores of these 47 animals in relation to the larger superset, we found that they had a similar distribution, spanning approximately +/- 1 std from the mean of the larger cohort (Figure 1B). Behavioral data was analyzed using scripts written in python 3.6 (Python Software Foundation, https://www.python.org/) using IPython (Granger, 2007) that utilized pandas (McKinney, 2010), scipy (Eric Jones, 2001), and statsmodels (Seabold, 2010) modules.

### Cortisol Assessment

Plasma samples were assayed for cortisol in duplicate using the DPC Coat-a-count radioimmunoassay (Siemens, Los Angeles, CA). The inter-assay CV%s calculated for a low and a high control were based on 21 assays. The low control had an average value of 48.4 µg/dL and a CV% of 7.0 and the high control had an average value of 128.3 µg/dL and a CV% of 7.0. The limit of detection defined by the lowest standard was 1 µg/dL.

### Tissue Collection for RNA Sequencing

For RNA-Seq blood was collected by femoral venipuncture into PreAnalytiX PAX gene RNA tubes (VWR, Radnor, PA). To collect brains, animals were euthanized under deep anesthesia with the guidance of veterinary staff using pentobarbital, which is the standard method of humane euthanasia at these facilities. This method of euthanasia is consistent with the recommendations of the Panel on Euthanasia of the American Veterinary Medical Association. Fresh frozen tissue was collected, cut into slabs, flash frozen in 2-methylbutane, and stored at −80°C as previously described (Fox et al., 2012). One hemisphere of the brain was cut into 14mm slabs.

### Laser Capture Microdissection Methods

#### Tissue preparation

Brain slabs used for LCM were counterbalanced for hemisphere across animals. The slab containing the CeL was selected for each animal. Acetylcholinesterase (AChE) staining was used to confirm that the slab contained the amygdala. In cases in which the amygdala was contained in two slabs both were used. Fourteen μm tissue sections were obtained on a cryostat at −20°C. Sections throughout the A-P extent of the CeL were used for LCM, mounted on Leica PEN 2.0µm membrane, laser capture microdissection slides (11532918, Leica, Wetzlar, Germany). Adjacent sections were used for AChE staining to localize the Ce and match the same level of CeL section across subjects (Figure 2A).

#### Rapid staining protocol

All staining was completed in an RNA-free environment and all steps were done on ice unless otherwise specified. LCM slides were removed from the −80°C freezer and thawed on a metal block for 30 seconds. Slides were fixed in acetone for 3 minutes at −20°C and then washed in PBS with 0.1% Triton x-100 (PBS-T) for 2 minutes. The tissue was stained with a neuron specific protein (NeuN) antibody (MAB377, Millipore, Burlington, MA) at a 1:200 dilution for 25 minutes at room temperature. Slides were submerged in PBS-T for 2 minutes and incubated in an HRP secondary antibody (PI-2000, Vectorlabs, Burlingame, CA) at 1:250 for 30 minutes at room temperature. Slides were then washed in PBS-T for 2 minutes and dipped in regular PBS for 30 seconds. Tissue was incubated in ImmPACT VIP peroxidase substrate (SK-4605, Vectorlabs) for 7-10 minutes before washing in DEPC-treated H_2_O for 5 minutes. Finally, the slides were dipped into 95% ethanol briefly before being dehydrated for 10-20 seconds each in 95% ethanol, 100% ethanol, and xylene. Slides were dried in a fume hood for 5 minutes before being transported to the laser capture microscope. Neurons were dissected using a Leica LMD6500 laser capture microscope. A circle that was approximately 850 µm^2^ was drawn around each neuron. Neurons were identified at 20x magnification, dissected with the laser, and fell into the lid of a microfuge tube (PCR-05-C, Axygen, Corning, NY) that was filled with 40uL of lysis buffer. After all neurons from the slide were captured, an additional 10uL of lysis buffer was added to the microfuge tube. The tube was vortexed on high for 30 seconds before being placed on dry ice.

#### Microdissection

Approximately 500-600 neurons were sampled from 6-8 slides (approximately every 0.25mm) that ranged the anterior-posterior extent of the CeL of each animal. After cells were dissected with LCM, each LCM slide image was overlaid with its adjacent AChE slide image in Adobe Illustrator CC 2014 to confirm that cells were dissected from the CeL. Tubes containing neuronal RNA were only used if at least 80% of the cells captured were identified as coming from the CeL. From our collection, we established 3 categories: 100% CeL cells, 80-99% CeL cells, and <80% CeL cells. Those slides that were <80% CeL cells were not used for RNA-Seq. To determine the “CeL neuron accuracy” for each animal, we calculated the proportion of the number of slides that contained 80-99% of CeL neurons out of total slides used for RNA-Seq (X 80-99% slides / (X 80-99% slides + Y 100% slides); Supplementary Figure 2B). AChE images were assigned a bregma value that most closely matched that AchE image in the Paxinos Atlas (Paxinos et al., 2009). A “weighted average bregma” value was calculated for each animal by multiplying the bregma value for each slide by the number of cells captured for in that slide and taking an average (Supplementary Figure 2B). For half of the animals (n=29), RNA from one slide was extracted and run on an Agilent Bioanalyzer to determine 18S and 28S peaks and RNA integrity numbers (average = 4, std = 0.75).

### RNA extraction and RNA Sequencing Pipeline

CeL LCM dissected neurons were pooled across sections within each animal prior to RNA extraction. RNA from neurons was extracted using Qiagen RNeasy Plus micro kit (74034, Qiagen, Hilden, Germany) with the modification for also purifying microRNAs. RNA samples were sequenced by Dr. Knowles (University of Southern California, Los Angeles, CA). Samples were processed with NuGen RNA-Seq V2 kit (7102-32, NuGen, San Carlos, CA) for cDNA synthesis and NuGEN Ovation Rapid library kit (0319, 0320, NuGen) for library preparation. For sequencing, the Illumina Hiseq 2500 with regular rapid sequencing prep kit was used. RNA was sequenced and a mean of 949710 mapped reads was found across animals. Reads were mapped to the 7.8 version of the rhesus monkey genome (MacaM; (Zimin et al., 2014) annotated by Dr. Norgren (University of Nebraska Medical Center, Omaha, NE). Mapping was performed using Sequence Alignment for Gene Expression (https://github.com/tadesouaiaia/sage) written in python2.7.

### RNA Sequencing Analysis and Statistics

#### Model Evaluation

To determine the most appropriate model for calculating differentially expressed (DE) genes, we first examined several regression models using ordinary least squares regression (OLS). The final model was also used for a confirmatory DE analysis in DESeq2. Genes that had one read in at least 20% of the animals were used for quantile normalization of the data and log2 transformed for use in OLS regression. Our goal was to build a model that assessed the association between gene expression and the predictor of interest. AT, freezing, cooing, and cortisol measured closest to time of death were used as predictors in individual models. We tested whether multiple factors that could contribute to variance and statistical inflation (Supplementary Table 1) should be included in our model. The statistical model was built within a framework designed to both maximize power to detect predictor-related associations while also reducing false positives discovered with permutation testing. We focused on models which described the largest fraction of the variance (R^2^) across the transcriptome without a cost to Bayesian Information Criterion (BIC; overfitting).

First, we explored each individual covariate in relation to gene expression across the transcriptome to identify potential candidates for the model. Next, we tested for collinearity among the potential covariates as well as our predictors of interest (Supplementary Figure 1A). Based on this, two measures (CeL neuron accuracy, age at necropsy (Nx)) were selected for model testing as they were not multicollinear and were the two variables with the greatest number of positive genes relative to false positives in a simulation (Supplementary Table 1). Sex was not selected for model testing because on its own it did not improve our ability to identify positive genes relative to false positives in a simulation as compared to other variables tested (Supplementary Table 1). Because sex was not included as a covariate, genes on chromosomes X and Y were removed to ensure that sex chromosome-related genes were also excluded from the analysis. Using CeL neuron accuracy and age at Nx as covariates separately as well as together, we tested models to establish the extent to which they could detect predictor-related gene expression relationships above chance. In the models that included the covariates, the predictor was shuffled and correlated with gene expression and this process was repeated 100 times to create a chance distribution. We chose the model that had the lowest BIC, accounted for the greatest percentage of the variance, and had the fewest false positives in a simulated model with a shuffled predictor. We settled on a final model which uses the covariates CeL neuron accuracy (see LCM methods: microdissection; Supplementary Table 1), and age at necropsy (Supplementary Table 1). This model was then used to create a more accurate chance distribution by running 10,000 simulations for each predictor of interest (Figure 4A). Analyses were focused on fully annotated genes where at least 50% of the samples expressed more than 1 mapped read.

#### Differential Gene Expression Analysis

The final model was used to determine differentially expressed (DE) genes that were associated with our predictor of interest. For each gene, we performed a permutation test to determine the probability of getting the nominal p-value for that gene by chance. This permutation test was separate from that used to construct the null distribution and evaluate the model. OLS regression and permutation testing were performed in python 2.7. Confirmatory analysis of DE genes was performed using DESeq2 (Love et al., 2014). Data were filtered and outliers removed based on a cook’s D distance of 3.8 which was calculated using a quantile function (qf) distribution as recommended in the DESeq2 protocol (Love et al., 2014). Genes that were DE in both analyses were the primary focus of further analyses. Gene ontologies were investigated using Panther to determine sets of AT-related genes that are associated with different molecular processes. For the heatmap (Figure 3A), the ward method of unsupervised clustering was performed in scripts written in python 3.6 (Python Software Foundation, https://www.python.org/) using the seaborn (version 0.0.9, https://seaborn.pydata.org) module which utilizes matplotlib (Hunter, 2007). Gene expression values in the heatmap are presented as log2 transformed, min-max scaled values. Gene expression values in scatterplots are quantile normalized, log2 transformed, and residualized for covariates.

### Immunohistochemistry: Tissue collection, PKCδ antibody characterization, staining protocols, and stereological analysis

To characterize the PKCδ antibodies, a brain from one additional monkey and brains from a wildtype (WT; C57BL/6J; stock 000664, Jackson Laboratories, Farmington, CT) and a PKCδ knockout (KO; B6;129X1-*Prkcd^tm1Msg^*/J; stock 028055, Jackson Laboratories) mouse were used. After characterization, perfused brains from three monkeys and three mice were used for between species comparison of PKCδ in the CeL. Animals were perfused using a standard protocol (Kalin et al., 2016). Brains were extracted, fixed overnight in 4% paraformaldehyde, cryoprotected in 30% sucrose, sectioned at 40-50μm and stored in cryoprotectant solution (30% ethylene glycol and 30% sucrose in 0.1 M phosphate buffer) at −20 °C (deCampo and Fudge, 2013).

### PKCδ Antibody Characterization for use in Rhesus Monkey Tissue

Three PKCδ antibodies (Supplementary Table 2) were assessed by staining both WT and PKCδ KO tissue using a standard immunohistochemistry protocol (see below). Those antibodies with specific binding to the PKCδ protein were used in subsequent immunofluorescence experiments (Supplementary Figure 2). The same antibodies were used to test for PKCδ staining in the monkey. To characterize specificity of PKCδ in monkey tissue, each antibody was preabsorbed with PKCδ peptide. Ten-fold molar excess of the PKCδ purified peptide was added to each antibody at their respective dilutions (Supplementary Table 2). The antibody-antigen mixture incubated for 36 hours at 4°C with mild agitation. The mixture was spun at 16,000g for 30 minutes. The resulting supernatant was pipetted into a clean vial and used in place of the primary antibody in the immunohistochemistry protocol.

### Immunohistochemistry protocol

Tissue from one mouse and monkey were stained together and all tissue was stained using the same lot of antibodies (Supplementary Table 2) and blocking serums. All tissue was removed from cryoprotectant solution 24 hours before staining and rinsed at least 3 times in 1x PBS. Primary antibodies were diluted in 1x PBS with 0.3% Triton x-100 and tissue was incubated overnight at room temperature. All secondary antibodies were made in 5% donkey serum. Fluorescent antibodies were also filtered through a 0.45μm filter to improve background staining.

Antibody characterization experiments were completed first and all staining was visualized using 3,3’-Diaminobenzidine (DAB). Tissue was incubated in 5% donkey serum (017-000-121, Jackson ImmunoResearch Laboratories, West Grove, PA) for 1 hour at room temperature. Tissue was subsequently washed three times in 1x PBS before being placed in PKCδ antibody overnight at room temperature. The following day, tissue for the antibody characterization experiments was washed in 1x PBS and incubated in an HRP conjugated secondary antibody diluted with blocking serum at 1:250 for 1 hour at room temperature. Tissue was then washed and incubated with ImmunoPact DAB (SK-4105, Vector Laboratories) for visualization of staining. Tissue was then mounted, left to dry overnight and then dehydrated in 75%, 95%, 100% ethanol solutions and xylene before being cover slipped. Images were captured at 1.25x, 5x and 20x magnification on a Leica LMD6500 microscope.

For triple labeling immunofluorescence, tissue was incubated serially in each antibody to avoid background. The staining protocols for the tissue used in the NeuN, PKCδ, and SST triple labeling experiment and the tissue used in the retrograde tracing experiment were the same. For the stereology experiment, tissue was first incubated in NeuN antibody overnight at room temperature, followed by 1x PBS washes and AlexaFluor 647 donkey anti-mouse secondary incubation for 1 hour at room temperature. For the retrograde tracing experiment, tissue was first incubated in anti-rhodamine, anti-WGA, or anti-fluorescein antibody overnight at room temperature, followed by 1x PBS washes and AlexaFluor 568 donkey anti-mouse secondary or AlexaFluor 488 donkey anti-goat secondary incubation for 1 hour at room temperature. Tissue was rinsed three times in 1x PBS and incubated in PKCδ antibody (1:200, Thermo Fisher) at room temperature for 24 hours for the mouse tissue and 48 hours for the monkey tissue. The following day tissue was washed three times in 1x PBS, incubated in AlexaFluor 488 donkey anti-rabbit for 1 hour at room temperature, washed three times in 1x PBS, and incubated in SST primary antibody for 72 hours to improve antibody penetration through the tissue. On the final day of staining, tissue was washed three times in 1x PBS and incubated in AlexaFluor 568 donkey anti-goat for 1 hour at room temperature before being washed in 1x PBS, incubated for 5 minutes in DAPI, mounted on slides, and coverslipped with Prolong Gold AntiFade Mountant (P36930, Thermo Fisher).

### Anatomical Boundaries of the Amygdala

The anatomical boundaries of the amygdala nuclei are well established in rodents (Price et al., 1987). We delineated the lateral division of the central nucleus in mouse tissue based on previously described work (Atlas, © 2018) and corroborated this with adjacent sections stained for AChE. In rhesus monkeys the nomenclature and anatomical boundaries have been previously described (Amaral et al., 1992; De Olmos, 2004). We used these descriptions to delineate the anatomical boundary for the lateral division of the central nucleus (CeL) in the rhesus monkey. We used the nomenclature system in the Paxinos atlas (Paxinos et al., 2009) for rhesus monkeys (Figure 5A) and for consistency, extend this nomenclature to the mouse.

### Imaging Protocol and Stereological Analysis

For the NeuN, PKCδ, and SST triple labeling experiment, the optical fractionator method was used to determine the total number of neurons, PKCδ+ neurons, SST+ neurons, and double labeled neurons in the CeL (Gundersen, 1986; West et al., 1991). Eight to ten sections per monkey and 4 sections per mouse through the anterior-posterior extent of the CeL was used and the first section was randomly selected within the first 6 (in mouse) or 8 (in monkey) sections. Each slide that contained the CeL was imaged with a 40x oil objective with a numerical aperture of 1.3 on a Nikon A1R+ confocal microscope. Stacks were acquired with a z-step of 1.25μm for monkey sections and 1μm for mouse sections and multiple stacks were stitched to produce whole CeL stacks to be used for offline stereological counting. Nikon nd2 files were exported from NIS elements into individual .tif files. Stereological counting was done in StereoInvestigator 2018.1.1 (MBF Bioscience, Williston, VT USA).

A sampling scheme was determined to estimate PKCδ and SST expressing neurons out of total neurons with an estimated Gunderson coefficient of error of 0.1 for each cell type. Neurons were counted when the center of the nucleolus of the NeuN stain came into focus. Cells expressing SST or PKCδ were counted only if they were also double labeled with NeuN. In a pilot study, NeuN and PKCδ+ neurons were counted at 25% in both species and SST+ and double labelled SST/PKCδ neurons were sampled at 100%. Using the sample-resample option in StereoInvestigator it was determined that in both species, PKCδ and NeuN labeled neurons could be sampled at a minimum of 2% and SST neurons could be sampled at a minimum of 10%. To ensure accurate counts, we chose to sample PKCδ and NeuN labeled cells at 5% and SST neurons at 20% in monkey tissue. SST/ PKCδ double labeled neurons were also sampled at 20% in monkey tissue and at 100% in mouse tissue. Section contours were determined based on CeL structure and NeuN staining and also followed the pattern visualized by adjacent AChE stained sections. For cell counting, section thickness was measured at every 10^th^ counting site. Counting frames were set at 75μm x 75μm in both mouse and monkey because frames of this size comfortably contained 4-6 cells. Tissue shrinkage was not consistently observed, and so guard zones were set at 5μm which still allowed for a 30μm disector height. To characterize the A-P distribution of the counted cells, cell counts were divided by the sampling percentage. Counts for each marker were then divided by the number of neurons counted in that section so that they could be presented as percentages.

Overall stereological estimates for each cell type are presented as percentage of the total number of neurons estimated. Between species differences for each cell type were tested using t-tests. Within species and for each cell type, an OLS regression model was used to investigate whether A-P location predicted number of cells counted. A variable for subject was included as a covariate in this model. A separate OLS model was run to test whether the A-P distribution for each cell type was different between species. One section from one monkey was not used as studentized residuals and cooks distance metrics demonstrated that it was an outlier. All statistical analyses were run in python 3.6 using the statsmodels (version 0.8.0) module.

### Surgery Procedure and Tissue Collection for Retrograde Injections

The surgical procedure has been previously described for these animals, which were used as part of other studies (deCampo and Fudge, 2013; Oler et al., 2017). The coordinates of the BST were localized for each animal prior to stereotaxic surgery by acquiring T2 weighted (structural) MRI images through the entire brain (3T, coronal sections, 0.8 mm thick, 0.1 mm apart). Animals were pre-anesthetized, intubated, and maintained on isofluorane throughout stereotaxic surgery for placement of injections. We placed multiple small injections (40 nL) of the bidirectional tracers, Lucifer yellow conjugated to dextran amine (LY; 10%, Molecular Probes, Eugene, OR), tetramethylrhodamine, conjugated to dextran amine (‘fluoruby’, FR; 4%, Molecular Probes), and fluorescein conjugated to dextran amine (FS; 10%, Molecular Probes) into the BSTL, in addition to several injections of the tracer wheat-germ agglutinin-horse radish peroxidase (WGA; 10%, Sigma, St. Louis, MO). Control injections of all tracers were placed in the nearby striatum. Previous studies from our laboratory have indicated that there is no cross-reactivity of antibodies to FR, FS, WGA and LY, permitting injections using different tracers into the same animal.

Two weeks after surgery, animals were deeply anesthetized and killed by perfusion through the heart with 0.9% saline containing 0.5 ml of heparin sulfate (200 ml/min for 10 minutes), followed by cold 4% paraformaldehyde in a 0.1 M phosphate buffer/30% sucrose solution (100 ml/min for 1 h). The brain was extracted from the skull, placed in a fixative overnight, and then put through increasing gradients of sucrose (10%, 20%, 30%). Brains were cut on a freezing microtome (40 μm) and all sections were stored in cryoprotectant solution (30% ethylene glycol and 30% sucrose in 0.1 M phosphate buffer) at −20 °C (Rosene et al., 1986). After confirmation of the injection and retrograde labeling using permanent immunostaining methods (Oler et al., 2017), additional sections through the extended amygdala for each case were selected for triple-labeling experiments.

### Retrograde Tracing Imaging Protocol and Analysis

Each slide that contained the CeL was investigated for the density of retrograde-labeled cells. For cases where there were many retrograde-labeled cells, the whole CeL was imaged and multiple stacks were stitched to produce whole CeL stacks to be used for offline counting. In cases where sections only expressed a small number of retrograde-labeled cells, individual stacks centered on the labeled cells were imaged and used for offline counting. Imaging parameters and image handling matched those used above. Retrograde-labeled neurons were first counted with channels for all other markers turned off. Once retrograde-labeled neurons were counted, each channel for the other marker was added to determine co-expression. Cells identified in the top 5μm and bottom 5μm were not counted. The overlap between retrograde-labeled neurons, PKCδ, SST, and SST/ PKCδ are presented as percentages. Venn diagrams were constructed with python 3.6 using the Venn3 module.

---

**Supplementary Figure 1.**
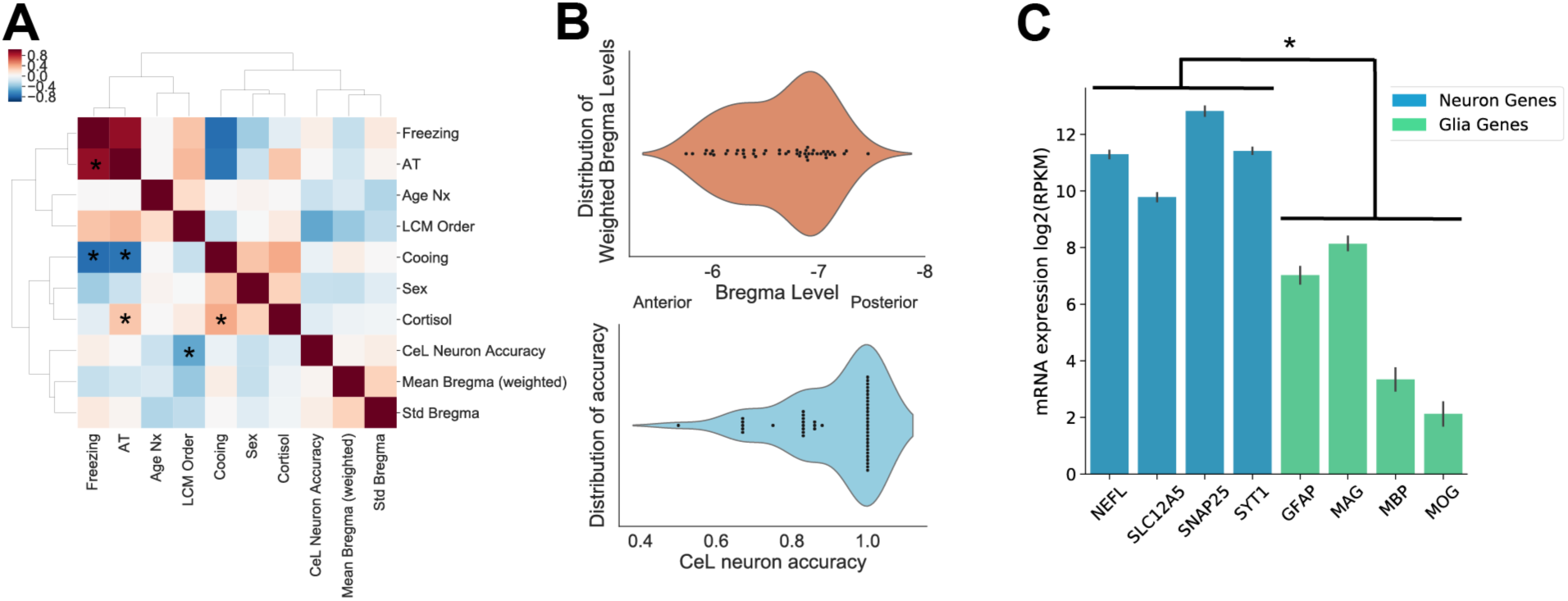
Characterization of LCM sampling method. (A) Heatmap depicting the Pearson correlations between model predictors (AT, freezing, cooing, and cortisol) as well as variables that were tested as covariates. Red indicates a positive correlation. Starred boxes indicate statistically significant Šidák-corrected p-values. (B) Top: Distribution of the weighted average bregma for each animal used for LCM, calculated by multiplying the bregma value of each slide by the number of cells captured in that slide and taking an average for each animal. Bottom: Distribution of CeL neuron accuracy for each animal, calculated as the proportion of the number of slides that contained 80-99% of CeL neurons out of total slides used for RNA-Seq for that animal. (C) Reads per kilobase million (RPKM) normalized mRNA expression values for neuron (blue) and glia (green) associated genes. Neuronal LCM samples expressed on average a greater amount of neuron-specific gene expression than glia-specific gene expression (p<0.0001, t=36.4; t-test). Error bars are displayed as SEM.

**Supplementary Figure 2.**
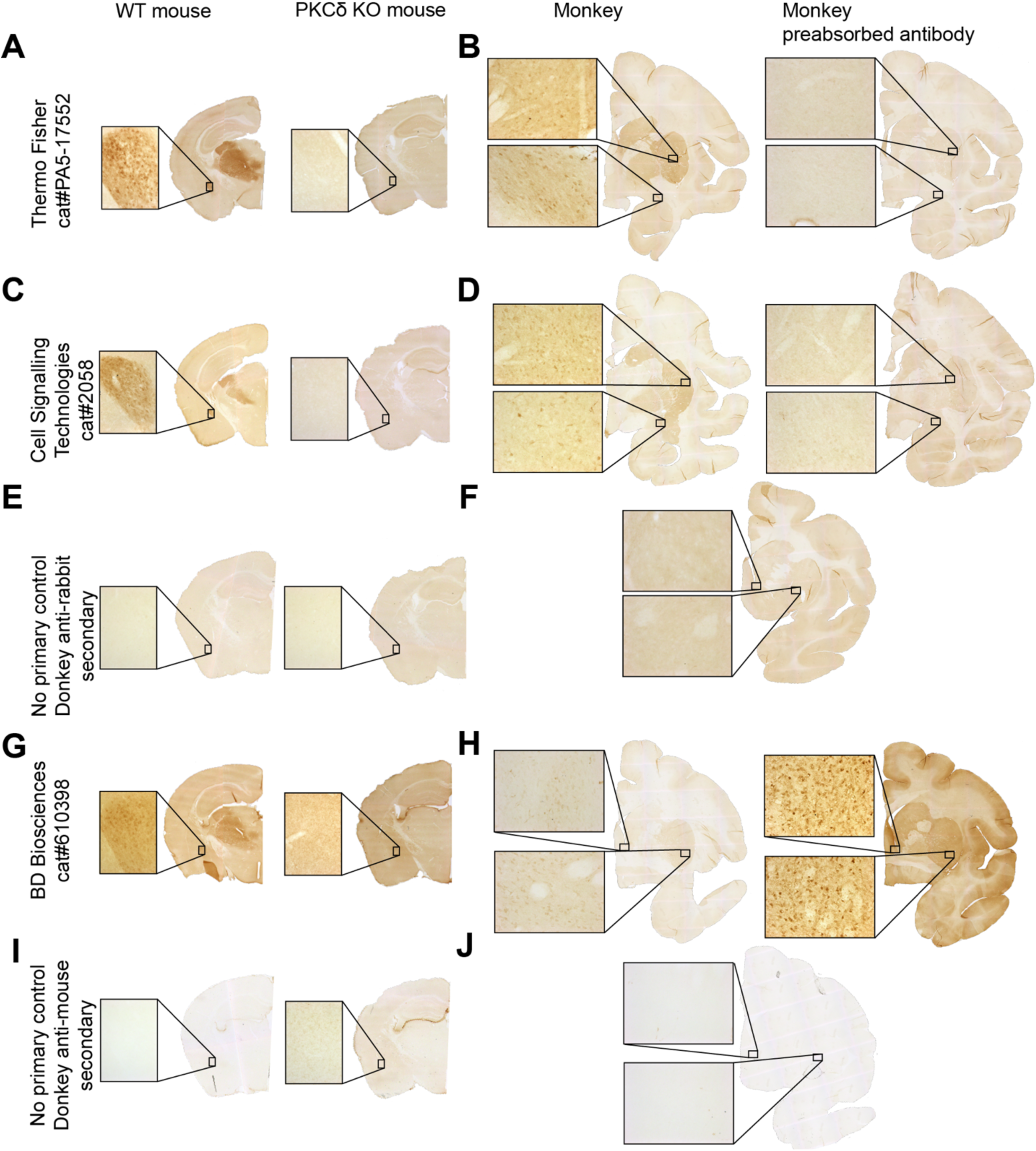
Characterization of PKCδ antibodies. Columns: Wild Type Mouse tissue, PKCδ Knock Out Tissue, Monkey tissue stained with PKCδ antibody, Monkey tissue stained with preabsorbed antibody. Rows: (A-B) ThermoFisher anti-PKCδ antibody, (C-D) Cell Signaling Technology anti-PKCδ antibody, (E-F) Donkey anti-rabbit secondary only control, (G-H) BD Biosciences anti-PKCδ antibody, (I-J) Donkey anti-mouse secondary only control. ThermoFisher and Cell Signaling Technology anti-PKCδ antibodies gave the most robust staining in both monkey and mouse tissue.

**Supplementary Figure 3.**
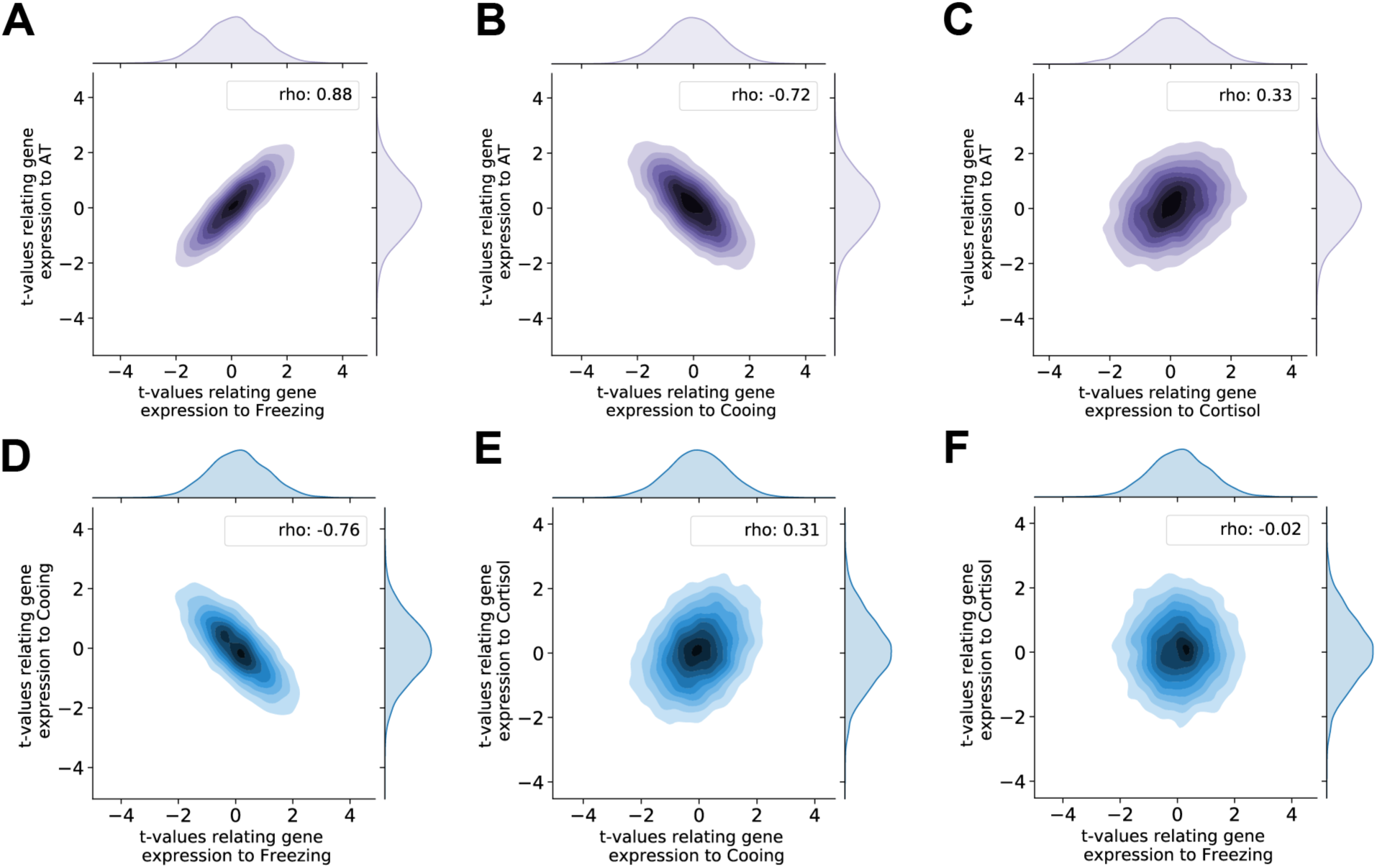
AT components share a molecular substrate. Kernel density plots depicting the distribution of t-values and Spearman correlations between t-values relating gene expression to (A) AT and freezing (rho=0.88, p<0.001) (B) AT and cooing (rho=,-0.72 p<0.001) (C) AT and cortisol (rho=0.33, p<0.001) (D) cooing and freezing (rho=-0.76, p<0.001) (E) cortisol and cooing (rho=0.31, p<0.001) (F) cortisol and freezing (rho=-0.02, p=0.27). Reported p-values are Šidák corrected.

**Supplementary Figure 4.**
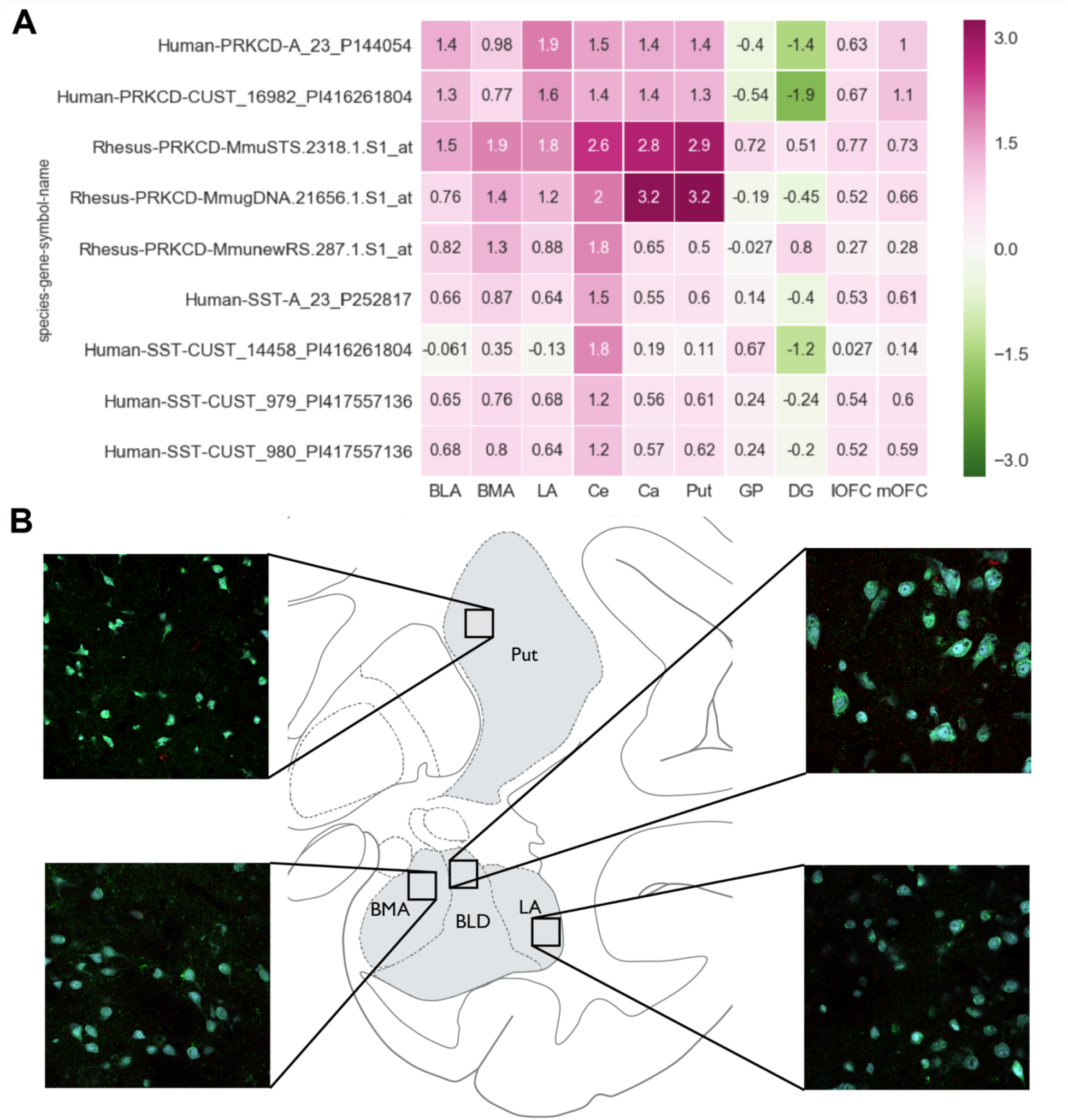
*PRKCD* mRNA and protein is widely distributed throughout the primate brain. (A) A heatmap of the human and nonhuman primate microarray data from the Allen Brain Atlas for *PRKCD* and *SST* demonstrate that the mRNA for these markers are widely distributed throughout multiple brain regions. Particular note is made of high levels of *PRKCD* in the Ce and striatum. The y-axis lists the different microarray probes for each marker for human and nonhuman primate. The x-axis describes the regions of the brain investigated. The color bar depicts the expression values, calculated using a z-score normalization of the relative expression level of each marker as compared to the rest of the brain (B). Immunofluorescence staining of PKCδ in the putamen as well as other regions of the nonhuman primate amygdala. BLD: dorsolateral division of the basal nucleus of the amygdala, BMA: basomedial nucleus of the amygdala, LA: lateral nucleus of the amygdala, Ce: central nucleus of the amygdala, Ca = caudate, put = putamen, GP: globus pallidus, DG: dentate gyrus of the hippocampus, lOFC: lateral orbitofrontal cortex, mOFC: medial orbitofrontal cortex.

## References

Ahrens, S., Wu, M.V., Furlan, A., Hwang, G.R., Paik, R., Li, H., Penzo, M.A., Tollkuhn, J., and Li, B. (2018). A Central Extended Amygdala Circuit That Modulates Anxiety. J Neurosci 38, 5567–5583.

Alheid, G.F., and Heimer, L. (1988). New perspectives in basal forebrain organization of special relevance for neuropsychiatric disorders: the striatopallidal, amygdaloid, and corticopetal components of substantia innominata. Neuroscience 27, 1–39.

Amador-Arjona, A., Elliott, J., Miller, A., Ginbey, A., Pazour, G.J., Enikolopov, G., Roberts, A.J., and Terskikh, A.V. (2011). Primary cilia regulate proliferation of amplifying progenitors in adult hippocampus: implications for learning and memory. J Neurosci 31, 9933–9944.

Amano, T., Amir, A., Goswami, S., and Pare, D. (2012). Morphology, PKCdelta expression, and synaptic responsiveness of different types of rat central lateral amygdala neurons. J Neurophysiol 108, 3196–3205.

Amaral, D.G., Avendano, C., and Benoit, R. (1989). Distribution of somatostatin-like immunoreactivity in the monkey amygdala. J Comp Neurol 284, 294–313.

Amaral, D.G., Price, J.L., Pitkanen, A., and Carmichael, S.T. (1992). Anatomical organization of the primate amygdaloid complex. In The Amygdala: Neurobiological Aspects of Emotion, Memory, and Mental Dysfunction (Wiley-Liss, Inc.), pp. 1–66.

Asok, A., Draper, A., Hoffman, A.F., Schulkin, J., Lupica, C.R., and Rosen, J.B. (2018). Optogenetic silencing of a corticotropin-releasing factor pathway from the central amygdala to the bed nucleus of the stria terminalis disrupts sustained fear. Molecular psychiatry 23, 914–922.

Atlas, A.B. (© 2018). Allen Brain Atlas API. (Allen Institute for Brain Science).

Berbari, N.F., Malarkey, E.B., Yazdi, S.M., McNair, A.D., Kippe, J.M., Croyle, M.J., Kraft, T.W., and Yoder, B.K. (2014). Hippocampal and cortical primary cilia are required for aversive memory in mice. PLoS ONE 9, e106576.

Biederman, J., Hirshfeld-Becker, D.R., Rosenbaum, J.F., Herot, C., Friedman, D., Snidman, N., Kagan, J., and Faraone, S.V. (2001). Further evidence of association between behavioral inhibition and social anxiety in children. Am J Psychiatry 158, 1673–1679.

Cai, H., Haubensak, W., Anthony, T.E., and Anderson, D.J. (2014). Central amygdala PKC-delta(+) neurons mediate the influence of multiple anorexigenic signals. Nat Neurosci 17, 1240–1248.

Carvajal, C., Dumont, Y., Herzog, H., and Quirion, R. (2006). Emotional behavior in aged neuropeptide Y (NPY) Y2 knockout mice. Journal of molecular neuroscience: MN 28, 239–245.

Cassel, M.D., and Gray, T.S. (1989). Morphology of Peptide Immunoreactive neurons in the rat central nucleus of the amygdala. Journal of Comparative Neurology, 320–333.

Cassell, M.D., Gray, T.S., and Kiss, J.Z. (1986). Neuronal architecture in the rat central nucleus of the amygdala: a cytological, hodological, and immunocytochemical study. J Comp Neurol 246, 478–499.

Chronis-Tuscano, A., Degnan, K.A., Pine, D.S., Perez-Edgar, K., Henderson, H.A., Diaz, Y., and al., e. (2009). Stable early maternal report of behavioral inhibition predicts lifetime social anxiety disorder in adolescence. J Am Acad Child Adolesc Psychiatry 48, 928–935.

Ciocchi, S., Herry, C., Grenier, F., Wolff, S.B., Letzkus, J.J., Vlachos, I., Ehrlich, I., Sprengel, R., Deisseroth, K., Stadler, M.B., et al. (2010). Encoding of conditioned fear in central amygdala inhibitory circuits. Nature 468, 277–282.

Cui, Y., Lv, G., Jin, S., Peng, J., Yuan, J., He, X., Gong, H., Xu, F., Xu, T., and Li, H. (2017). A Central Amygdala-Substantia Innominata Neural Circuitry Encodes Aversive Reinforcement Signals. Cell reports 21, 1770–1782.

Davidson, R.J., and Rickman, M. (1999). Behavioral inhibition and the emotional circuitry of the brain: Stability and plasticity during the early childhood years. In Extreme fear, shyness, and social phobia: Origins, biological mechanisms, and clinical outcomes, L.A. Schmidt, and J. Schulkin, eds. (NY: Oxford University Press), pp. 67–87.

De Olmos, J.S. (2004). The Amygdala. In The Human Nervous System Paxinos G., and Mai J.K., eds. (San Diego: Elsevier Academic Press).

deCampo, D.M., and Fudge, J.L. (2013). Amygdala projections to the lateral bed nucleus of the stria terminalis in the macaque: comparison with ventral striatal afferents. J Comp Neurol 521, 3191–3216.

Dong, H.W., Petrovich, G.D., and Swanson, L.W. (2001). Topography of projections from amygdala to bed nuclei of the stria terminalis. Brain research Brain research reviews 38, 192–246.

Duman, R.S., and Li, N. (2012). A neurotrophic hypothesis of depression: role of synaptogenesis in the actions of NMDA receptor antagonists. Philosophical transactions of the Royal Society of London Series B, Biological sciences 367, 2475–2484.

Eley, T.C., Bolton, D., O’Connor, T.G., Perrin, S., Smith, P., and Plomin, R. (2003). A twin study of anxiety-related behaviours in pre-school children. J Child Psychol Psychiatry 44, 945–960.

Eric Jones, T.O., Pearu Peterson, others (2001). SciPy: Open source scientific tools for Python.

Essex, M.J., Klein, M.H., Slattery, M.J., Goldsmith, H.H., and Kalin, N.H. (2010). Early risk factors and developmental pathways to chronic high inhibition and social anxiety disorder in adolescence. Am J Psychiatry 167, 40–46.

Fadok, J.P., Krabbe, S., Markovic, M., Courtin, J., Xu, C., Massi, L., Botta, P., Bylund, K., Muller, C., Kovacevic, A., et al. (2017). A competitive inhibitory circuit for selection of active and passive fear responses. Nature 542, 96–100.

Fadok, J.P., Markovic, M., Tovote, P., and Luthi, A. (2018). New perspectives on central amygdala function. Current opinion in neurobiology 49, 141–147.

Fox, A.S., and Kalin, N.H. (2014). A translational neuroscience approach to understanding the development of social anxiety disorder and its pathophysiology. Am J Psychiatry 171, 1162–1173.

Fox, A.S., Oler, J.A., Shackman, A.J., Shelton, S.E., Raveendran, M., McKay, D.R., Converse, A.K., Alexander, A., Davidson, R.J., Blangero, J., et al. (2015a). Intergenerational neural mediators of early-life anxious temperament. Proc Natl Acad Sci U S A 112, 9118–9122.

Fox, A.S., Oler, J.A., Shelton, S.E., Nanda, S.A., Davidson, R.J., Roseboom, P.H., and Kalin, N.H. (2012). Central amygdala nucleus (Ce) gene expression linked to increased trait-like Ce metabolism and anxious temperament in young primates. Proc Natl Acad Sci U S A 109, 18108–18113.

Fox, A.S., Oler, J.A., Tromp do, P.M., Fudge, J.L., and Kalin, N.H. (2015b). Extending the amygdala in theories of threat processing. Trends Neurosci 38, 319–329.

Fox, A.S., Shelton, S.E., Oakes, T.R., Converse, A.K., Davidson, R.J., and Kalin, N.H. (2010). Orbitofrontal cortex lesions alter anxiety-related activity in the primate bed nucleus of stria terminalis. Journal of Neuroscience 30, 7023–7027.

Fox, A.S., Shelton, S.E., Oakes, T.R., Davidson, R.J., and Kalin, N.H. (2008). Trait-like brain activity during adolescence predicts anxious temperament in primates. PLoS ONE 3, e2570.

Fox, A.S., Souaiaia, T., Oler, J.A., Kovner, R., Kim, J.M.H., Nguyen, J., French, D.A., Riedel, M.K., Fekete, E.M., Rabska, M.R., et al. (2019). Dorsal Amygdala Neurotrophin-3 Decreases Anxious Temperament in Primates. Biol Psychiatry.

Fox, N.A., Henderson, H.A., Marshall, P.J., Nichols, K.E., and Ghera, M.M. (2005). Behavioral inhibition: linking biology and behavior within a developmental framework. Annual review of psychology 56, 235–262.

Frank, S. (2009). Tetrabenazine as anti-chorea therapy in Huntington disease: an open-label continuation study. Huntington Study Group/TETRA-HD Investigators. BMC neurology 9, 62.

Frank, S. (2010). Tetrabenazine: the first approved drug for the treatment of chorea in US patients with Huntington disease. Neuropsychiatric disease and treatment 6, 657–665.

Granger, F.P.a.B.E. (2007). IPython: A System for Interactive Scientific Computing. Computing in Science & Engineering 9, 21–29.

Gray, T.S., Cassell, M.D., and Williams, T.H. (1982). Synaptology of three peptidergic neuron types in the central nucleus of the rat amygdala. Peptides 3, 273–281.

Gray, T.S., and Magnuson, D.J. (1992). Peptide immunoreactive neurons in the amygdala and the bed nucleus of the stria terminalis project to the midbrain central gray in the rat. Peptides 13, 451–460.

Greene, M.W., Morrice, N., Garofalo, R.S., and Roth, R.A. (2004). Modulation of human insulin receptor substrate-1 tyrosine phosphorylation by protein kinase Cdelta. The Biochemical journal 378, 105–116.

Guay, D.R. (2010). Tetrabenazine, a monoamine-depleting drug used in the treatment of hyperkinetic movement disorders. The American journal of geriatric pharmacotherapy 8, 331–373.

Gundersen, H.J. (1986). Stereology of arbitrary particles. A review of unbiased number and size estimators and the presentation of some new ones, in memory of William R. Thompson. Journal of microscopy 143, 3–45.

Gungor, N.Z., Yamamoto, R., and Pare, D. (2015). Optogenetic study of the projections from the bed nucleus of the stria terminalis to the central amygdala. J Neurophysiol 114, 2903–2911.

Han, W., Tellez, L.A., Rangel, M.J., Jr., Motta, S.C., Zhang, X., Perez, I.O., Canteras, N.S., Shammah-Lagnado, S.J., van den Pol, A.N., and de Araujo, I.E. (2017). Integrated Control of Predatory Hunting by the Central Nucleus of the Amygdala. Cell 168, 311–324.e318.

Haubensak, W., Kunwar, P.S., Cai, H., Ciocchi, S., Wall, N.R., Ponnusamy, R., Biag, J., Dong, H.W., Deisseroth, K., Callaway, E.M., et al. (2010). Genetic dissection of an amygdala microcircuit that gates conditioned fear. Nature 468, 270–276.

Hettema, J.M., Neale, M.C., and Kendler, K.S. (2001). A review and meta-analysis of the genetic epidemiology of anxiety disorders. Am J Psychiatry 158, 1568–1578.

Hirshfeld, D.R., Rosenbaum, J.F., Biederman, J., Bolduc, E.A., Faraone, S.V., Snidman, N., Reznick, J.S., and Kagan, J. (1992). Stable behavioral inhibition and its association with anxiety disorder. J Am Acad Child Adolesc Psychiatry 31, 103–111.

Hunter, J.D. (2007). Matplotlib: A 2D Graphics Environment. Computing in Science & Engineering 9, 90–95.

Janak, P.H., and Tye, K.M. (2015). From circuits to behaviour in the amygdala. Nature 517, 284–292.

Kagan, J., Reznick, J.S., and Snidman, N. (1987). The physiology and psychology of behavioral inhibition in children. Child Dev 58, 1459–1473.

Kalin, N.H., Fox, A.S., Kovner, R., Riedel, M.K., Fekete, E.M., Roseboom, P.H., Tromp do, P.M., Grabow, B.P., Olsen, M.E., Brodsky, E.K., et al. (2016). Overexpressing Corticotropin-Releasing Factor in the Primate Amygdala Increases Anxious Temperament and Alters Its Neural Circuit. Biol Psychiatry 80, 345–355.

Kalin, N.H., and Shelton, S.E. (1989). Defensive behaviors in infant rhesus monkeys: environmental cues and neurochemical regulation. Science 243, 1718–1721.

Kalin, N.H., and Shelton, S.E. (2003). Nonhuman Primate Models to Study Anxiety, Emotion Regulation, and Psychopathology. Annals of the New York Academy of Sciences 1008, 189–200.

Kalin, N.H., Shelton, S.E., and Davidson, R.J. (2004). The role of the central nucleus of the amygdala in mediating fear and anxiety in the primate. J Neurosci 24, 5506–5515.

Kalin, N.H., Shelton, S.E., Davidson, R.J., and Kelley, A.E. (2001). The primate amygdala mediates acute fear but not the behavioral and physiological components of anxious temperament. Journal of Neuroscience 21, 2067–2074.

Kalin, N.H., Shelton, S.E., Fox, A.S., Oakes, T.R., and Davidson, R.J. (2005). Brain regions associated with the expression and contextual regulation of anxiety in primates. Biol Psychiatry 58, 796–804.

Kanehisa, M., and Goto, S. (2000). KEGG: kyoto encyclopedia of genes and genomes. Nucleic Acids Res 28, 27–30.

Kanehisa, M., and Sato, Y. (2019). KEGG Mapper for inferring cellular functions from protein sequences. Protein Sci.

Kim, J., Zhang, X., Muralidhar, S., LeBlanc, S.A., and Tonegawa, S. (2017). Basolateral to Central Amygdala Neural Circuits for Appetitive Behaviors. Neuron 93, 1464–1479.e1465.

Kim, S.Y., Adhikari, A., Lee, S.Y., Marshel, J.H., Kim, C.K., Mallory, C.S., Lo, M., Pak, S., Mattis, J., Lim, B.K., et al. (2013). Diverging neural pathways assemble a behavioural state from separable features in anxiety. Nature 496, 219–223.

Kovner, R., Fox, A.S., French, D.A., Roseboom, P.H., Oler, J.A., Fudge, J.L., and Kalin, N.H. (2019). Somatostatin Gene and Protein Expression in the Non-human Primate Central Extended Amygdala. Neuroscience 400, 157–168.

LeDoux, J.E., Cicchetti, P., Xagoraris, A., and Romanski, L.M. (1990). The lateral amygdaloid nucleus: sensory interface of the amygdala in fear conditioning. J Neurosci 10, 1062–1069.

Li, H., Penzo, M.A., Taniguchi, H., Kopec, C.D., Huang, Z.J., and Li, B. (2013). Experience-dependent modification of a central amygdala fear circuit. Nat Neurosci 16, 332–339.

Li, L.X., Zhou, J.X., Calvet, J.P., Godwin, A.K., Jensen, R.A., and Li, X. (2018). Lysine methyltransferase SMYD2 promotes triple negative breast cancer progression. Cell Death & Disease 9, 326.

Love, M.I., Huber, W., and Anders, S. (2014). Moderated estimation of fold change and dispersion for RNA-seq data with DESeq2. Genome Biology 15, 550.

Lubieniecka, J.M., de Bruijn, D.R.H., Su, L., van Dijk, A.H.A., Subramanian, S., van de Rijn, M., Poulin, N., van Kessel, A.G., and Nielsen, T.O. (2008). Histone Deacetylase Inhibitors Reverse SS18-SSX–Mediated Polycomb Silencing of the Tumor Suppressor *Early Growth Response 1* in Synovial Sarcoma. Cancer Research 68, 4303–4310.

Martin, L.J., Powers, R.E., Dellovade, T.L., and Price, D.L. (1991). The bed nucleus-amygdala continuum in human and monkey. J Comp Neurol 309, 445–485.

McCullough, K.M., Morrison, F.G., Hartmann, J., Carlezon, W.A., Jr., and Ressler, K.J. (2018). Quantified Coexpression Analysis of Central Amygdala Subpopulations. eNeuro 5.

McKinney, W. (2010). Data Structures for Statistical Computing in Python. In Proceedings of the 9th Python in Science Conference, pp. 51–56.

Moga, M.M., and Gray, T.S. (1985). Evidence for corticotropin-releasing factor, neurotensin, and somatostatin in the neural pathway from the central nucleus of the amygdala to the parabrachial nucleus. J Comp Neurol 241, 275–284.

Munoz-Estrada, J., Lora-Castellanos, A., Meza, I., Alarcon Elizalde, S., and Benitez-King, G. (2018). Primary cilia formation is diminished in schizophrenia and bipolar disorder: A possible marker for these psychiatric diseases. Schizophrenia research 195, 412–420.

Newton, A.C. (2010). Protein kinase C: poised to signal. Am J Physiol Endocrinol Metab 298, E395–402.

Oler, J.A., Fox, A.S., Shelton, S.E., Rogers, J., Dyer, T.D., Davidson, R.J., Shelledy, W., Oakes, T.R., Blangero, J., and Kalin, N.H. (2010). Amygdalar and hippocampal substrates of anxious temperament differ in their heritability. Nature 466, 864–868.

Oler, J.A., Fox, A.S., Shackman, A.J., and Kalin, N.H. (2016). The central nucleus of the amygdala is a critical substrate for individual differences in anxiety. In Living without an Amygdala. In In Living without an Amygdala (New York, NY: Guilford Press), pp. 218–251.

Oler, J.A., Tromp, D.P., Fox, A.S., Kovner, R., Davidson, R.J., Alexander, A.L., McFarlin, D.R., Birn, R.M., B, E.B., deCampo, D.M., et al. (2017). Connectivity between the central nucleus of the amygdala and the bed nucleus of the stria terminalis in the non-human primate: neuronal tract tracing and developmental neuroimaging studies. Brain Struct Funct 222, 21–39.

Pare, D., and Smith, Y. (1993). The intercalated cell masses project to the central and medial nuclei of the amygdala in cats. Neuroscience 57, 1077–1090.

Paxinos, G., Huang, X., Petrides, M., and Toga, A.W. (2009). The Rhesus Monkey Brain: Stereotaxic Coordinates (Elsevier).

Petrovich, G.D., and Swanson, L.W. (1997). Projections from the lateral part of the central amygdalar nucleus to the postulated fear conditioning circuit. Brain Res 763, 247–254.

Pomrenze, M.B., Tovar-Diaz, J., Blasio, A., Maiya, R., Giovanetti, S.M., Lei, K., Morikawa, H., Hopf, F.W., and Messing, R.O. (2019). A Corticotropin Releasing Factor Network in the Extended Amygdala for Anxiety. J Neurosci 39, 1030–1043.

Price, J.L., Russchen, F.T., and Amaral, D.G. (1987). The limbic region. II. The amygdaloid complex. In Handbook of Chemical Neuroanatomy, B.T. Hokfelt, and L.W. Swanson, eds. (Amsterdam: Elsevier), pp. 279–381.

Pruski, M., and Lang, B. (2019). Primary Cilia-An Underexplored Topic in Major Mental Illness. Frontiers in psychiatry 10, 104.

Rankin, S.L., Guy, C.S., Rahimtula, M., and Mearow, K.M. (2008). Neurotrophin-induced upregulation of p75NTR via a protein kinase C-delta-dependent mechanism. Brain Res 1217, 10–24.

Roberts, G.W., Woodhams, P.L., Polak, J.M., and Crow, T.J. (1982). Distribution of neuropeptides in the limbic system of the rat: the amygdaloid complex. Neuroscience 7, 99–131.

Roseboom, P.H., Nanda, S.A., Fox, A.S., Oler, J.A., Shackman, A.J., Shelton, S.E., Davidson, R.J., and Kalin, N.H. (2014). Neuropeptide Y receptor gene expression in the primate amygdala predicts anxious temperament and brain metabolism. Biol Psychiatry 76, 850–857.

Rosene, D.L., Roy, N.J., and Davis, B.J. (1986). A cryoprotection method that facilitates cutting frozen sections of whole monkey brains for histological and histochemical processing without freezing artifact. J Histochem Cytochem 34, 1301–1315.

Sajdyk, T.J., Schober, D.A., Smiley, D.L., and Gehlert, D.R. (2002). Neuropeptide Y-Y2 receptors mediate anxiety in the amygdala. Pharmacology, biochemistry, and behavior 71, 419–423.

Sawyers, C., Ollendick, T., Brotman, M.A., Pine, D.S., Leibenluft, E., Carney, D.M., Roberson-Nay, R., and Hettema, J.M. (2019). The genetic and environmental structure of fear and anxiety in juvenile twins. American journal of medical genetics Part B, Neuropsychiatric genetics: the official publication of the International Society of Psychiatric Genetics 180, 204–212.

Schindelin, J., Arganda-Carreras, I., Frise, E., Kaynig, V., Longair, M., Pietzsch, T., Preibisch, S., Rueden, C., Saalfeld, S., Schmid, B., et al. (2012). Fiji: an open-source platform for biological-image analysis. Nature Methods 9, 676.

Seabold, S., and Josef Perktold (2010). Statsmodels: Econometric and statistical modeling with python.

Shackman, A.J., Fox, A.S., Oler, J.A., Shelton, S.E., Davidson, R.J., and Kalin, N.H. (2013). Neural mechanisms underlying heterogeneity in the presentation of anxious temperament. Proc Natl Acad Sci U S A 110, 6145–6150.

Stein, M.B., Jang, K.L., and Livesley, W.J. (1999). Heritability of anxiety sensitivity: a twin study. Am J Psychiatry 156, 246–251.

Tang, L., Nogales, E., and Ciferri, C. (2010). Structure and Function of SWI/SNF Chromatin Remodeling Complexes and Mechanistic Implications for Transcription. Progress in biophysics and molecular biology 102, 122–128.

Tasan, R.O., Nguyen, N.K., Weger, S., Sartori, S.B., Singewald, N., Heilbronn, R., Herzog, H., and Sperk, G. (2010). The central and basolateral amygdala are critical sites of neuropeptide Y/Y2 receptor-mediated regulation of anxiety and depression. J Neurosci 30, 6282–6290.

Taylor, T.N., Caudle, W.M., Shepherd, K.R., Noorian, A., Jackson, C.R., Iuvone, P.M., Weinshenker, D., Greene, J.G., and Miller, G.W. (2009). Nonmotor symptoms of Parkinson’s disease revealed in an animal model with reduced monoamine storage capacity. J Neurosci 29, 8103–8113.

Tschenett, A., Singewald, N., Carli, M., Balducci, C., Salchner, P., Vezzani, A., Herzog, H., and Sperk, G. (2003). Reduced anxiety and improved stress coping ability in mice lacking NPY-Y2 receptors. Eur J Neurosci 18, 143–148.

Turner, C.A., Clinton, S.M., Thompson, R.C., Watson, S.J., Jr., and Akil, H. (2011). Fibroblast growth factor-2 (FGF2) augmentation early in life alters hippocampal development and rescues the anxiety phenotype in vulnerable animals. Proc Natl Acad Sci U S A 108, 8021–8025.

Tye, K.M., Prakash, R., Kim, S.Y., Fenno, L.E., Grosenick, L., Zarabi, H., Thompson, K.R., Gradinaru, V., Ramakrishnan, C., and Deisseroth, K. (2011). Amygdala circuitry mediating reversible and bidirectional control of anxiety. Nature 471, 358–362.

Veening, J.G., Swanson, L.W., and Sawchenko, P.E. (1984). The organization of projections from the central nucleus of the amygdala to brainstem sites involved in central autonomic regulation: a combined retrograde transport-immunohistochemical study. Brain Res 303, 337–357.

Viviani, D., Charlet, A., van den Burg, E., Robinet, C., Hurni, N., Abatis, M., Magara, F., and Stoop, R. (2011). Oxytocin selectively gates fear responses through distinct outputs from the central amygdala. Science 333, 104–107.

Wang, W.J., Tay, H.G., Soni, R., Perumal, G.S., Goll, M.G., Macaluso, F.P., Asara, J.M., Amack, J.D., and Tsou, M.F. (2013). CEP162 is an axoneme-recognition protein promoting ciliary transition zone assembly at the cilia base. Nature cell biology 15, 591–601.

Wang, Z., Phan, T., and Storm, D.R. (2011). The type 3 adenylyl cyclase is required for novel object learning and extinction of contextual memory: role of cAMP signaling in primary cilia. J Neurosci 31, 5557–5561.

West, M.J., Slomianka, L., and Gundersen, H.J. (1991). Unbiased stereological estimation of the total number of neurons in thesubdivisions of the rat hippocampus using the optical fractionator. The Anatomical record 231, 482–497.

Ye, J., and Veinante, P. (2019). Cell-type specific parallel circuits in the bed nucleus of the stria terminalis and the central nucleus of the amygdala of the mouse. Brain Struct Funct.

Yu, K., Ahrens, S., Zhang, X., Schiff, H., Ramakrishnan, C., Fenno, L., Deisseroth, K., Zhao, F., Luo, M.H., Gong, L., et al. (2017). The central amygdala controls learning in the lateral amygdala. Nat Neurosci 20, 1680–1685.

Yu, K., Garcia da Silva, P., Albeanu, D.F., and Li, B. (2016). Central Amygdala Somatostatin Neurons Gate Passive and Active Defensive Behaviors. J Neurosci 36, 6488–6496.

Zimin, A.V., Cornish, A.S., Maudhoo, M.D., Gibbs, R.M., Zhang, X., Pandey, S., Meehan, D.T., Wipfler, K., Bosinger, S.E., Johnson, Z.P., et al. (2014). A new rhesus macaque assembly and annotation for next-generation sequencing analyses. Biology Direct 9, 20.

